# Activity-Based Protein Profiling Identifies *Klebsiella pneumoniae* Serine Hydrolases with Potential Roles in Host-Pathogen Interactions

**DOI:** 10.1101/2024.05.28.596221

**Authors:** Md Jalal Uddin, George Randall, Jiyun Zhu, Tulsi Upadhyay, Laura van Eijk, Paul B. Stege, Frerich M. Masson, Marco C. Viveen, Matthew Bogyo, Matthias Fellner, Marcel R. de Zoete, Mona Johannessen, Christian S. Lentz

## Abstract

*Klebsiella pneumoniae* is a normal resident of the human gastro-intestinal tract and an opportunistic, critical priority pathogen that can cause a variety of severe systemic infections. Due to emerging multi-drug resistance of this pathogen, the discovery and validation of novel targets for the development of new treatment options is an urgent priority. Here, we explored the family of serine hydrolases, a highly druggable and functionally diverse enzyme family which is uncharacterized in *K. pneumoniae*. Using functionalized covalent fluorophosphonate inhibitors as activity-based probes we identified 10 serine hydrolases by mass spectrometry-based activity-based protein profiling, 7 of which were previously uncharacterized. Functional validation using transposon mutants deficient in either of the putative lysophospholipase PldB, esterase YjfP and patatin-like phospholipase YchK revealed severe growth defects in human colonic organoid co-culture models and reduced virulence during *Galleria mellonella* infection. Mutants deficient in the PldB and YjfP, but not YchK show increased susceptibility to killing by complement and the antimicrobial peptide antibiotic polymyxin B, suggesting a role in maintaining cell envelope integrity. Biochemical characterization and structural analysis of recombinant YjfP suggest this protein is a deacetylase. This study gives important insights into the molecular mechanisms underlying virulence and cell physiology of *K. pneumoniae* at the host-pathogen interface and it positions PldB, YjfP and YchK as potential antimicrobial or anti-virulence target candidates, inhibition of which might synergize with existing antibiotics and human immune defenses.

## Introduction

The Gram-negative enterobacterium *Klebsiella pneumoniae* is an opportunistic pathogen that naturally resides in the gastrointestinal tract of healthy humans and animals but can cause a variety of severe extra-intestinal infections, including urinary tract, bloodstream, and lung infections^1^. This bacterium is also known for its ability to produce extended-spectrum beta-lactamases (ESBLs) and carbapenemases enzymes that confer resistance to many antibiotics, including the carbapenems, which are often used as last-resort antibiotics ^2,3^. This makes the infections caused by *K. pneumoniae* difficult to treat and increases the risk of mortality ^4^. Carbapenemase-producing *K. pneumoniae* are critical priority pathogens on the list of multi-drug resistant (MDR) pathogens by the World Health Organization (WHO). Therefore, research and development of new therapeutics need to be prioritized to prevent entry into a post-antibiotic era^5^ Beyond multi-drug resistance the emergence and global spread of hypervirulent *K. pneumoniae* strains, which are common causes of liver abscesses and other systemic infections are of great concern ^6,7^.

In the light of the spread of multi-drug resistant strains of this pathogen, it is essential to understand the molecular factors that contribute to its pathogenicity and validate their potential as drug targets. One particularly attractive family of putative target enzyme are the serine hydrolases (SHs), a large and diverse enzyme class with roles in various cellular processes, from metabolism, to signaling, and regulation of gene expression, and can be putative drug targets for a variety of diseases ^8–10^. Due to their conserved mechanism and active site architecture, many SHs share reactivity towards the same active site-directed inhibitors, such as fluorophosphonates. This can be exploited by a chemoproteomic technique called activity-based protein profiling (ABPP) that uses functionally-tagged active site-directed covalent enzyme inhibitors (activity-based probes, ABPs) to detect active enzyme species under physiological conditions in complex samples of interest^11–14^. Fluorescent fluorophosphonate (FP)-probes can be used to detect SHs in gel-based ABPP, whereas FP-biotin allows for pull-down and identification of targets by liquid chromatography-mass spectrometry (LC-MS) (Schematically illustrated in **Fig. 1A, B**). In this way, SH activities have been profiled in diverse biological samples including animal tissues^15^, human cell lines ^16^, and more recently bacterial pathogens including *Staphylococcus aureus* ^17^, *S. epidermidis* ^18^, *Mycobacterium tuberculosis* ^19–21^, and *Vibrio cholerae* ^22^, the gut commensal *Bacteroides thetaiotaomicron*^23^ and archaea ^24^. In our recent chemoproteomic study on *S. aureus*, we identified a family of ten largely uncharacterized fluorophosphonate-binding serine hydrolases FphA-J several of which have roles in pathogenesis ^17^ and bacterial stress responses ^25^. These enzymes can be targeted by specific small molecule inhibitors and probes holding promise as anti-virulence ^17^ and pathogen-specific imaging targets^26^.

**Figure 1.**
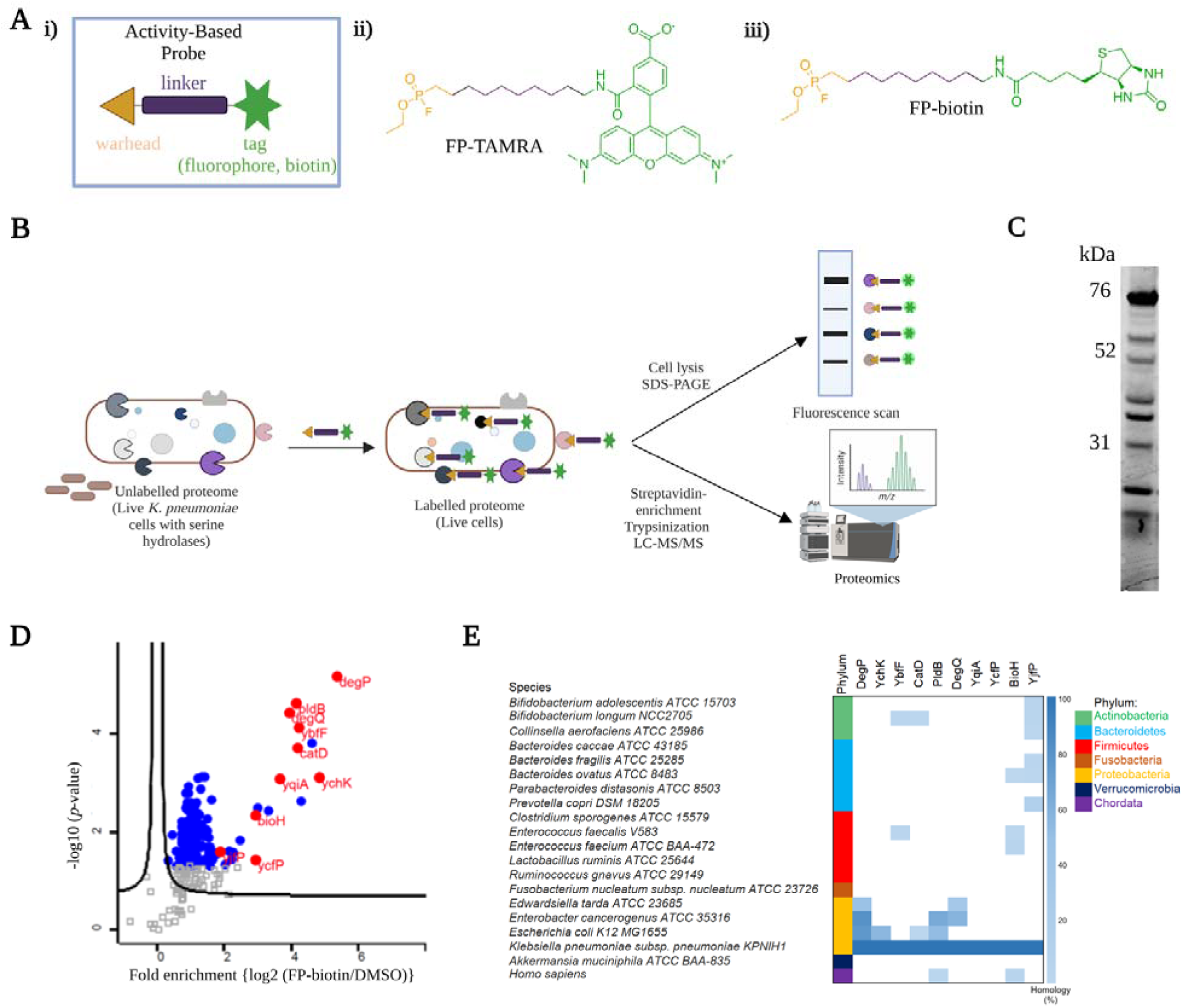
Chemoproteomic identification of serine hydrolases in *K. pneumoniae*. **A**) Schematic drawing of the features of Activity-based probes (ABPs) (i) and the chemical structures of the ABPs FP-TAMRA (ii) and FP-bioti (iii) that were used in this study. **B**) Schematic overview of general workflow for activity-based protein profiling (ABPP) of serine hydrolases in live *K. pneumoniae*. Fluorophosphonate-based ABPs selectively bind to accessible and active serine hydrolase species. Fluorescently tagged ABPs allow their visualization by SDS-PAGE analysis and fluorescence scanning, whereas biotinylated probes are used for streptavidin-enrichment and identification via LC-MS/MS or they can be visualized using a fluorescent tag through SDS-PAGE and in-gel fluorescence analysis. **C)** SDS-PAGE analysis of *K. pneumoniae* MKP103 live cells labeled with FP-TMR for 60 min at 37°C. The graph depicts fluorescent scans in the Cy3 (520nm) channel utilizing the Amersham™ Typhoon™ 5 (cytiva) imagin system. **D)** Volcano plot of *K. pneumoniae* proteins identified by LC-MS/MS that were enriched after FP-bioti treatment compared to a vehicle-treated control dataset. A two-tailed two-sample t-test was conducted to compare cells labeled with DMSO and FP-biotin. Significantly enriched hits are highlighted with blue dots, significantly enriched 10 SHs are indicated with red dots, and enriched but not significant hits are marked with grey rectangles. **E)** Heatmap of homologs of *K. pneumoniae* serine hydrolases across 20 representative gut commensal bacterial species and in humans (Chordata), as sourced from the Human Microbiome Project Reference Genomes for the Gastrointestinal Tract database using BLAST-P. In the heatmap, each filled cell indicates that the species has a homolog of the *K. pneumoniae* SHs, as determined by a threshold e-value of 1 × 10^−10^; white background denotes the absence of a homolog.

Surprisingly, very few serine hydrolases of *K. pneumoniae* have been described or functionally characterized. A mutant of the periplasmic HtrA-like serine protease DegP showed higher susceptibility to complement-mediated killing and reduced levels of capsule polysaccharide ^27^. However, it remains unclear how this protease mediates these effects. In *E. coli*, the two HtrA-serine proteases DegP and DegS have described functions in extracytoplasmic protein quality control and stress responses (reviewed in^28^).

Another serine hydrolase, the phospholipase Tle1 has been described as a secreted effector of the type 6 secretion system (T6SS) in a hypervirulent *K. pneumoniae* strain^29,30^. Due to toxic effects of T6SS effectors on both bacterial competitors and host tissues, the *K. pneumoniae* T6SS is important for both long-term colonization of the gut^30^ as well as intestinal barrier dysfunction and bacterial translocation from the gut^31^.

Here, we performed a systematic analysis of serine hydrolase activities in *K. pneumoniae* using a broad-spectrum serine hydrolase probe. Seven out of 10 identified hydrolases were poorly annotated and previously uncharacterized. Functional characterization using transposon mutants in colonization/infection assays involving HT29-MTX cells, human-derived 2D colonic organoids, and *Galleria mellonella* infection suggest crucial roles for at least three of these hydrolases during infection and/or colonization. Our data suggest that the cell membrane-associated putative lysophospholipase PldB and esterase YjfP contribute to cell envelope integrity and contribute to resistance to antimicrobial peptides and complement, whereas the putatively secreted patatin-like phospholipase YchK may contribute to *K. pneumoniae* virulence by degrading mucus or host-cell derived phospholipids. Biochemical and structural analysis of recombinant YjfP suggest that this enzyme functions as a deacetylase.

## Materials and Methods

### Bacterial strains and culture conditions

Strains of *K. pneumoniae* MKP103 and its isogenic mutants are summarized in **Supplementary Table 1**. All strains were routinely cultured on Blood agar or Luria-Bertani broth (LB broth). The bacterial strains were incubated at 37°C, and liquid cultures were aerated by shaking at 180 rpm, except when intended for co-incubation with mammalian cells or organoids (described in their respective sections below).

### Bacterial labeling with fluorophosphonate-tetramethylrhodamine (FP-TAMRA)

After overnight growth on either an agar plate or in liquid culture, as indicated, the bacteria were adjusted to the desired density in LB/MH broth and added to microtubes in a final volume of 50-100 μL. ActivX™ TAMRA-FP (ThermoFisher Scientific) (at a concentration of 1 μM) was added from 100X stock solutions in DMSO and the cells were incubated for 60 minutes at 37 °C and 300 RPM. Following probe labeling, bacterial suspensions were transferred to 2 mL screw-cap tubes filled with 30-50 µL of 4x SDS-Loading buffer (comprising 40% glycerol, 240 mM Tris/HCl at pH 6.8, 8% SDS, 0.04% bromophenol blue, and 5% beta-mercaptoethanol) and 60-100 µL of 0.1 mm glass beads, and then lysed using bead-beating (3×30s, 6500 rpm, with 60s pause in-between) (Precellys® Evolution homogenizer (Bertin Technologies) and centrifuge samples at 6000g for 5 min at 4°C to remove the debris.

### SDS-PAGE analysis of fluorescently labeled proteins

After adding 4x SDS sample buffer, the samples were subjected to boiling at 95 °C for 10 minutes and subsequently separated using SDS-PAGE gel electrophoresis. The resulting gels were scanned for fluorescence scanning in the Cy3 (532nm) channel, utilizing the Amersham™ Typhoon™ 5 imaging system (cytiva).

### FP-biotin labeling of *K. pneumoniae* and sample preparation for mass-spectrometry

*K. pneumoniae* MKP103 cultures were grown on a blood plate for 24 hours and resuspended to an OD_600_ ∼20 in 3 mL MH broth. For each biological replicate, 1 mL aliquots were transferred to a 1.5 mL tube and either FP-Biotin (3 μM) or DMSO was added, and the cells were then incubated for 60 min at 37°C and 700 rpm before samples were centrifuged at 4,500 ×g for 5 minutes at 4°C, and the supernatant was removed. The cell pellets were resuspended in 1.2 mL RIPA Lysis buffer (50 mM Tris, 150 mM NaCl, 0.1% SDS, 0.5% sodium deoxycholate, 1% Triton X-100) in 2 mL screw-cap tube filled with ca. 200 µL of 0.1 mm glass beads and lysed by bead-beating (3×30s, 6500 rpm, with 60s pause in-between, performed two to three times with one min interval on ice). After centrifugation for 5 minutes at 10,000 ×g at 4°C, the protein concentration in the supernatant was adjusted to 1.0 mg/mL, and the proteins were stored at –20°C until the sample preparation.

For each sample, 50 μL of streptavidin magnetic beads were washed twice with 1 mL of RIPA lysis buffer. The streptavidin beads were then incubated with 1 mg of protein from each sample in an additional 500 μL of RIPA lysis buffer at 4°C overnight on a rotator set at 18 RPM. After enrichment, the beads were collected using a magnetic rack and washed with RIPA lysis buffer twice (1 mL, 2 minutes at room temperature), followed by a wash with 1 M KCl (1 mL, 2 minutes at room temperature), a wash with 0.1 M Na_2_CO_3_ (1 mL, ∼10 seconds), a wash with 2 M urea in 10 mM Tris-HCl (pH 8.0) (1 mL, ∼10 seconds), and two washes with RIPA lysis buffer (1 mL per wash, 2 minutes at room temperature). After the final wash, the beads were transferred to fresh protein Lo-Bind tubes with 1 mL of RIPA lysis buffer and washed three times with 500 µL of 4M Urea in 50 mM Ammonium bicarbonate (Ambic) with shaking for 7 minutes each time to remove nonspecific enrichments. Lastly, the beads were washed three times with 500 µL of 50 mM Ambic with shaking for 7 minutes, changing the tube between these washes.

For on-bead digestion, 150 μL of 50mM Ambic, 3 μL of 1mM CaCl2, 0.75 μL of 1M DDT, 4.5 μL of 500mM IAA, and 6 μL of MS-grade trypsin solution were added to the protein Lo-Bind tube, and samples were incubated at 37 °C overnight with a shaker running at 800 rpm. After digestion, tryptic peptide digests were separated, and beads were washed with 70 µL of 50 mM Ambic. For each sample, 20 μL of formic acid was added to the combined eluates. The samples were stored at –20°C until LC-MS/MS analysis. The sample preparation was conducted using the same method previously described^32^.

### Liquid chromatography–mass spectrometry analysis

Sample cleanup and concentration were performed using OMIX C18 tip (A5700310, Varian). For the LC-MS analysis, the concentrated samples were dissolved in 15 µl of 0.1% formic acid. Then, 0.5 µg of peptides from each sample were injected for analysis. Peptide mixtures containing 0.1% formic acid were loaded onto EASY-nLC1200 system (Thermo Fisher Scientific) with a C18 column (2 µm, 100 Å, 50 µm, 50 cm), and fractionation was performed using a 5-80% acetonitrile gradient in 0.1% formic acid at a flow rate of 300 nL/min for 60 minutes. The fractionated peptides were analyzed using a Orbitrap Exploris 480 mass spectrometer (Thermo Fisher Scientific). Data acquisition was carried out in a data-dependent mode employing a Top20 method. Raw data was processed using Proteome Discoverer 3.1 software with the CHIMERYS, and the fragmentation spectra were matched against the (*Klebsiella pneumoniae* subsp. pneumoniae KPNIH1, Taxonomy ID: 1087440) database. A peptide mass tolerance of 10 ppm and a fragment mass tolerance of 0.02 Da were employed during the search. Peptide ions were filtered using a false discovery rate (FDR) set at 5% for accurate peptide identifications. To ensure precision at least three biological replicates were conducted for all samples. Statistical analysis was conducted using Perseus software (version 2.03.0)^33^. Potential contaminants, reverse hits, and proteins identified by side only were excluded. The intensities from label-free quantification were converted using a log2 transformation. We applied imputation based on a normal distribution (width, 0.3; down-shift, 1.8) to handle missing values. The p-values were determined through a two-sided, two-sample t-test. Proteins identified as significantly enriched serine hydrolases had at least a threefold change (equivalent to a log2 fold-change of 1.58) and a minimum P value of 0.05 (corresponding to a −log10 value of 1.30). The mass spectrometry proteomics data have been deposited to the ProteomeXchange Consortium via the PRIDE^34^ partner repository with the dataset identifier PXD052404.

### Bioinformatic analyses

The 10 putative serine hydrolases were analyzed using both uniport^35^ and the web-based InterPro tool (https://www.ebi.ac.uk/interpro/) to predict proteins structures. The subcellular localization of serine hydrolases in *K. pneumoniae* was predicted with the web-based tools PSORTb v3.0 (https://www.psort.org/psortb/index.html)^36^. Furthermore, homology of *K. pneumoniae* serine hydrolases was identified by querying the full-length protein sequences against the non-redundant protein sequences database for gut commensal bacteria and *Homo sapiens* using Blastp with an E value cutoff of 10^-10^ for all identified homologs.

### Growth analyses

Growth curves were monitored using 96-well microtiter plates. Overnight cultures of *K. pneumoniae* MKP103 WT and its transposon mutants were diluted 1:100 in fresh medium or in medium supplemented with polymyxin B at a concentration of 8 µg/mL. A 200 µL aliquot of these diluted cultures was transferred to each well of the 96-well plate as the initial culture. The plates were then incubated at 37°C, and the optical density at 600 nm (OD600) was measured every 10 minutes using a Synergy H1 Hybrid Reader (BioTek) or a Bioscreen plate reader. The area under growth curve (AUC) was determined by using GraphPad Prism 9 for polymyxin B treatment.

### Co-incubation of HT29-MTX cells and *K. pneumoniae*

The HT29-MTX human colorectal adenocarcinoma cell line ^37^ was grown in Dulbecco’s modified Eagle medium (DMEM, supplied by Thermo Fisher) supplemented with 10% heat-inactivated (HI) fetal calf serum (FCS). For coculture experiments, HT-29-MTX cells were seeded at a density of approximately 1 x 10^5^ cells per well in 48-well plates and mid-log phase *K. pneumoniae* MKP103 and SH-deficient mutants (with OD600 nm of 0.4) in FBS-free DMEM were added to each well at a multiplicity of infection (MOI) of 20. After centrifugation at 200 x g for 5 minutes, the plates were incubated at 37°C with 5% CO_2_ for 2, 4, 6, and 8 hours and samples were collected from the media above host cells for counting the colony-forming units per milliliter (CFU/mL). All experiments were performed in duplicate and repeated independently at least three times.

### Organoid line and growth conditions

The clonal human-derived colonic organoid cell line Pt15-70206 was used to grow colonic epithelium, following the protocol described by Vonk *et al*. ^38^. Briefly, colonic organoid stocks were thawed, and the organoids were cultured in Matrigel domes with medium containing 15% Advanced DMEM/F12, 1x Glutamax, 100 U/mL Penicillin-Streptomycin, 10 mmol/L HEPES (all Invitrogen), 25% Rspo1 Conditioned Medium, 10% Noggin Conditioned Medium, 0.5nM Wnt Surrogate-FC Fusion Protein (ImmunoPrecise-UCN001), 2% B27 (Invitrogen), 1.25 mM N-acetylcysteine, 10 mM Nicotinamide, 3μM p38 inhibitor SB202190 (all Sigma-Aldrich), 50 ng/mL mEGF (Invitrogen), and 0.5 μM A83-01 (Tocris).

After an average of 4 passages, the organoids were disrupted and used to seed transwells (corning, 3470-clear) to generate confluent colonic epithelium. During the first 24 hours in transwells, the organoid culture medium was supplemented with 10nM rock inhibitor Y-27632 (Sigma-Aldrich). Once the Trans-epithelial Electrical Resistance (TEER) surpassed 100 Ω/cm^2^, the culture medium was replaced with differentiation medium^39^, excluding Wnt Surrogate-Fc Fusion Protein, nicotinamide, and p38 inhibitor. Differentiation medium was refreshed 24 hours before the adding bacteria for co-culturing, in absence of Penicillin-Streptomycin.

### Co-incubation of organoids and *K. pneumoniae*

Overnight *K. pneumoniae* MKP103 and its SH-deficient transposon mutants were diluted 1/100 in fresh medium and cultured in LB at 37°C until reaching the exponential growth phase with an optical density at 600nm of 0.3-0.4. Subsequently, the bacterial suspension was washed in PBS and resuspended in PBS to an OD600 of 0.4. The bacterial suspension was further diluted in organoid medium to achieve∼2.5× 10^6^ CFU/mL when adding them to the 2D organoid monolayers. The transwells containing the organoids and bacteria were spun down at 250 x g for 2 minutes to ensure cell contact. Finally, the co-culture was incubated at 37°C with 5% CO_2_ for 2, 4, 6, 8, and 16 hours and samples were collected from the apical side supernatant for counting the CFUs. To evaluate bacterial replication on the epithelial surface, the monolayer tissue and membrane were removed from the insert and dissociated using 2 mm glass beads in ice-cold PBS through 30 s of vortexing. Alternatively, after removal from the insert, the cells were lysed using 0.2% Triton X-100. The resulting PBS suspension was serially diluted and plated on LB agar. The agar plates were incubated overnight at 37°C, and CFUs were counted. In parallel, samples were taken from the basolateral compartment, serially diluted, and plated to assess any breach of the monolayer. The CFUs were manually enumerated and expressed as CFU/ml. The epithelial cells were examined for damage with the EVOS FL Auto 2 fluorescent microscopes (ThermoFisher Scientific) at 16 hours.

### Visualization of 2D organoid cultures

Organoid 2D cultures were fixed with Methacarn fixation (Methanol-based Carnoy fixation), which causes protein denaturation^40^, consisting of 60% methanol, 30% chloroform, and 10% glacial acetic acid, for two hours. After washing away the fixative with 100% methanol, the samples were rehydrated using different ethanol dilutions (100%, 90%, 80%, 70%, 60%, 40%, 20%), with each step lasting ten minutes. The filter of the transwells, with the organoids, was separated from the insert and blocked and permeabilized with a blocking buffer containing 2% Bovine Serum Albumin (BSA) (Serva) and 0.1% saponin (Sigma Aldrich) before incubating with the primary and secondary antibodies at 4°C in dark. The primary antibodies used were Anti-MUC1 21D4 (Sigma Aldrich, 05-652), Anti-MUC2 (Abcam, ab76774), Anti-MUC13 (generous gift by Karin Strijbis), Anti-Sox9 (Merck Millipore, AB5535), Anti-Villin 1D2C3 (Santa Cruz, sc-58897), and Anti-lysozyme (Dako, A099). Then, organoids were stained with Phalloidin Atto-647 (65906-10NMOL) and DAPI and imaged with an SP5 II confocal microscope (Leica TCS).

### Membrane permeabilization

Bacteria cultured overnight were diluted 1/100 in fresh medium and grown to an OD600 of 0.4–0.5 at 37°C with shaking. The bacteria were washed with RPMI with 0.05% human serum albumin (HSA, Sanquin, The Netherlands) through centrifugation at 10,000×g for 2 minutes and resuspended to an OD600 of 0.5 in RPMI. For the assay, bacteria were adjusted to an OD600 of 0.05 and mixed with 10 % normal human serum (NHS) (serum was prepared as described in^41^) in the presence of 1 µM SYTOX Green Nucleic Acid stain (ThermoFisher) in RPMI. This mixture was incubated at 37°C with shaking. Fluorescence was measured using a CLARIOstar microplate reader (Labtech) at an excitation wavelength of 490–14 nm and an emission wavelength of 537–30 nm.

### Protein Expression and Purification

The full length *yjfP*, *yqiA*, and *pldB* (Uniprot^35^ IDs A6THA0, A6TE20, and A6TGK5) DNA sequences were codon optimized for *E. coli* expression (IDT and TISIGNER^42^) and synthesized by IDT with overhangs for ligation-independent cloning^43^. The synthesized DNAs were cloned into a modified pET28a-LIC vector incorporating an N-terminal His_6_-tag and a 3C protease cleavage site. Plates were incubated at 37 °C overnight and single colonies were used to inoculate a starter culture of 5 mL LB supplemented with 50 µg/mL kanamycin. The starter culture was used to inoculate 1 L of LB media supplemented with 50 µg/mL kanamycin and incubated at 37°C while shaking at 180 rpm until the OD_600nm_ reached 0.5. The cultures were incubated at 18 °C while shaking at 180 rpm and recombinant protein expression was induced with 1 mM IPTG. Cell pellets were harvested the following day by centrifugation at 5,000 × g for 30 minutes. Cell pellets were either stored at – 20°C or lysed right away for protein purification.

A cell pellet was resuspended in lysis buffer (50 mM Tris pH 8.0, 300 mM NaCl, 50 mM imidazole, 10% sucrose, 10% glycerol) and incubated on ice for 10 minutes with 400 µg lysozyme and 200 µg DNase. The cells were subsequently lysed via sonication in an ice bath using Sonifier Heat Systems Ultrasonics, for 5 min using a one second pulse mode. Lysate was cleared by centrifugation at 15,000 x g for 30 minutes. The supernatant was loaded onto 2.0 mL Ni^2+^-NTA resin which was pre-washed with lysis buffer. The resin-supernatant mixture was incubated at 4 °C room for 30 min before the resin was washed in lysis buffer. The recombinant protein was eluted via 7 mL of elution buffer (50 mM Tris pH 8.0, 300 mM NaCl, 300 mM imidazole, 10% sucrose, 10% glycerol). For PldB and YjfP, these purified His_6_-tagged proteins were used for biochemical analysis described below. To cleave the N-terminal His_6_ tag for protein crystallization studies, the eluted fractions of YqiA and YjfP incubated with 5 mM DTT and 3C protease overnight at 4 °C. Recombinant proteins of YjfP and YqiA were finally purified by size exclusion chromatography (gel filtration). Protein samples were concentrated using 10,000 MWCO spin columns to approximately 250 µL and injected onto a size exclusion column (Superdex 75 Increase 10/300 GL column, GE Life Sciences) preequilibrated with size exclusion buffer (25 mM Tris pH 8.0, 150 mM NaCl). Protein purity was assessed by SDS-PAGE and nanodrop. Samples were concentrated to ∼10 mg/mL and snap frozen in liquid nitrogen or used immediately for crystallization.

### Biochemical assay of PldB, YjfP, and YqiA

Activity tests of purified serine hydrolases on different 4-methylumbelliferyl(MU) substrates were carried out as previously described^44^. Briefly, the reaction was carried out in PBS with 0.02 % v/v Triton X-100 (Fisher Scientific, Fairlawn, NJ). 8 µL enzyme solution (0.625 nM His_6_-PldB, 0.625 nM His_6_-YjfP, 62.5 nM YqiA (tag-free)) was mixed with 2 µL 5x 4-MU substrate stock solutions (250 µM) to initiate reactions and then the fluorescence signal (λ_ex_= 335 nm and λ_em_= 450 nm) was monitored by Cytation 3 Multi-Mode Reader (BioTek, Winooski, VT) at 25 °C for 1 hour. Turnover rates of the 4-MU substrates in the linear phase of the reaction were calculated in GraphPad Prism9 as RFU/second. The reaction rates were corrected by subtraction of the background hydrolysis in the absence of proteins for each substrate. The substrate preference profile was generated by dividing the corrected reaction rates by enzyme concentrations in each assay.

### Protein Crystallisation

YjfP and YqiA were crystallized using sitting drop vapour diffusion and screened for crystallization against several broad screens. 0.2 µL of protein solutions at 10 mg/mL were mixed with 0.2 µL of mother liquor. YjfP appeared as needle-like crystals in several different conditions, however the final structure of YjfP was determined from cubic crystals grown in 0.1 M ammonium sulfate, 35 % w/v PEG 8,000, 0.1 M sodium acetate pH 5.0. A YjfP crystal from this condition was soaked in mother liquor supplemented with 25 % v/v ethylene glycol for one minute before freezing in liquid nitrogen for analysis at the Australian Synchrotron. YqiA crystallized in several conditions that mostly contained either calcium chloride or calcium acetate, a crystal grown in 0.3 M calcium chloride hexahydrate, 10 % w/v PEG 6,000, 0.1 Tris pH 8.0 led to successful structure determination of YqiA. This crystal was soaked in mother liquor supplemented with 25 % v/v glycerol for one minute, before being frozen in liquid nitrogen.

### Crystal data collection and Processing

Protein crystals were subject to x-ray diffraction at the Australian Synchrotron MX2 beamline^9^. Datasets were processed in XDS^10^, before being merged and scaled using AIMLESS^11^ in the CCP4 suite. Phases were determined by molecular replacement in Phenix Phaser^12^ using an AlphaFold model^13^ of YqiA or YjfP prepared with Phenix Process Predicted Model^14^. The output model underwent iterative building and refinement using COOT^15^ and Phenix^16^.

### Infection of K. pneumoniae in G. mellonella larvae

*Galleria mellonella* larvae were obtained from Reptilutstyr AS (Norway). For infection experiments, *K. pneumoniae* MKP103 and its transposon mutants were cultured overnight in LB broth and harvested by centrifugation at 4,500 × g for 10 min at 4°C followed by one wash with PBS. The bacteria were then diluted in PBS to achieve OD600 of 1, approximately 1 × 10^9^ CFU/ml. Each experimental group consisted of n=10 larvae. For infection, 10 µl of a bacterial suspension adjusted to 1 × 10^5^ CFU/ml in PBS was injected into right hid proleg using a 30G syringe microapplicator (0.30 mm (30G) × 8 mm, BD Micro-Fine demi). A separate group of 10 larvae were injected with 10 µl of PBS to ensure that death was not due to injection trauma as negative control. The larvae were placed in 9.2 cm Petri dishes and incubated at 37°C in the dark and survival was monitored for 5 days. Each experiment was independently replicated at least three times.

### Statistical analysis

Graphs and statistical analyses were produced using GraphPad Prism 9. Unless specified otherwise, all experiments were conducted using three independent biological replicates. Data for graphs are shown as mean ± standard deviation (SD). To compare wild-type (WT) to mutant strains, either one-way or two-way ANOVA tests were utilized, followed by Dunnett’s multiple comparisons test, using the WT strain as the control. Significance levels were denoted as follows: *P < 0.05, **P < 0.01, ***P < 0.001, and ****P < 0.0001.

## Results

### Identification serine hydrolases in *K. pneumoniae*

We first set out to determine global serine hydrolase (SH) activity profile in *K. pneumoniae* when grown on blood agar using the fluorescent ABP FP-TMR. We selected strain MKP103, a derivative of the multi-drug resistant clinical outbreak strain KPNIH1 for which the carbapenemase-gene has been deleted^45^. KPNIH1 belongs to sequence type ST258 which can cause life-threatening infections in hospitalized individuals and poses a considerable risk to spread multidrug resistance into the community^46,47^. After labeling, the cells were lysed, and the labeled proteins were separated via SDS-PAGE analysis and visualized by in-gel fluorescence scanning. The results revealed that several bands were predominantly labeled at the 1 µM probe concentration (**Fig. 1C**). To identify these SH targets, we then used a biotiynylated probe (FP-biotin) with a chemoproteomic workflow using streptavidin-enrichment and LC-MS/MS analysis (**Fig. 1 Aiii, B**). Using this approach, we identified 10 *K. pneumoniae* SHs that displayed significant enrichment (p-value < 0.05, enrichment > 3.5-fold) when compared to a control dataset that was not treated with FP-biotin (as shown in **Fig. 1B. E** and detailed in **Supplementary Table 2** and **Extended Dataset 1,** data are available from ProteomeXchange with identifier PXD052404). Of the 10 identified SHs, 8 were predicted to possess α,β-hydrolase domains, while the remaining two were annotated as serine proteases with Trypsin-like peptidase domains (**Table 1**). Most of these enzymes are generally not well characterized in *K. pneumoniae* and for many of these enzymes their cellular functions are ill defined. Of the 10 SHs, only the HtrA-protease DegP and the carboxylesterase BioH enzymes have been studied functionally in *K. pneumoniae*. The latter putatively hydrolyze the pimeloyl-acyl carrier protein methyl ester as one of the first steps of biotin biosynthesis. ^48^ Sequence-based bioinformatic prediction of the subcellular location of the 10 identified SH proteins using PSORTb v3.0^36^ suggests that only the patatin-like phospholipase YchK is presumably secreted, whereas the two HtrA-like proteases, DegP and DegQ, were predicted to localize to the periplasmic. YjfP and YqiA were predicted to localize to the cytoplasmic. YbIF and YcfA are unknown. PldB is on the cytoplasmic membrane, and BioH and CatD to the cytosol. Except for DegP, DegQ, YchK and YqiA, most of the *K. pneumoniae* SHs displayed limited or no homology with serine hydrolases from 19 other gut commensals and human (**Fig. 1E, Supplementary Table 3, Extended Dataset 2**).

**Table 1.** Overview of serine hydrolases identified in *Klebsiella pneumoniae subsp. pneumoniae* KPNIH1. Refer to Supplementary Dataset **1** and Supplementary Table 2.

|  | **Gene name** | **Functional Annotation (putative) (Pfam)** | **Predicted location (Putative) (PSORTb)** | **% Homology with *H. sapiens* protein** | **MW [kDa]** |
| --- | --- | --- | --- | --- | --- |
| KPSH 1 | *degP* | Trypsin-like peptidase | Periplasmic | - | 49.5 |
| KPSH 2 | *ychK* | Patatin-like phospholipase | Extracellular | - | 33.3 |
| KPSH 3 | *ybfF* | AB_Hydrolase 6 | Unknown | - | 28.5 |
| KPSH 4 | *catD* | Hydrolase 4 | Cytoplasmic | - | 27.3 |
| KPSH 5 | *pldB* | Hydrolase 4 | Cytoplasmic membrane | 25% | 38.2 |
| KPSH 6 | *degQ* | Trypsin-like peptidase | Periplasmic | - | 47.2 |
| KPSH 7 | *yqiA* | UPF0227 | Cytoplasmic | - | 21.5 |
| KPSH 8 | *ycfP* | UPF0227 | Cytoplasmic | - | 21.1 |
| KPSH 9 | *bioH* | AB_Hydrolase 1 | Cytoplasmic | 30% | 28.3 |
| KPSH 10 | *yjfP* | Peptidase S9 | Unknown | - | 26.5 |

To functionally validate the identified target enzymes, we retrieved transposon mutant strains with insertions in individual serine hydrolase genes (*yfb*F*, yjf*P*, yqi*A*, ych*K*, deg*P*, pldB,* d*eg*Q, and *cat*D) from the Manoil Lab Transposon Mutant Library^45^. The library did not contain mutants in *bio*H or *ycf*P. At least for BioH with its presumable role in the biosynthesis of biotin, the absence of viable transposon mutants might indicate essentiality. We first labelled the obtained transposon mutants with FP-TMR labeled protein bands in the gel-based FP-TMR labeling. This allowed us to confirm the identities of *pldB*, *degP*, *ychK*, and *ybfF*, but not *degQ, yjfP*, and *ycfF*, as their corresponding transposon mutants showed similar FP-TMR profiles as the WT (**Fig. 2A**). This lack of labeling in the gel based ABPP assay suggesting that some of the identified enzymes may not be abundant and can only be detected in the more sensitive MS assay. In addition, five fluorescent protein bands could not be assigned to the MS identified serine hydrolases, suggesting that differences in selectivity or cellular permeability of the biotinylated and fluorescent probes might have prevented them from being detected in the proteomics approach (**Fig. 2B**). Based on this outcome, a set of five targets were selected for further functional studies: YfbF, YchK, PldB, DegP and the uncharacterized esterase YjfP which lacks homologs outside of the *Enterobacteriales* and humans (**Fig. 1E, Supplementary Table 3**)

**Figure 2.**
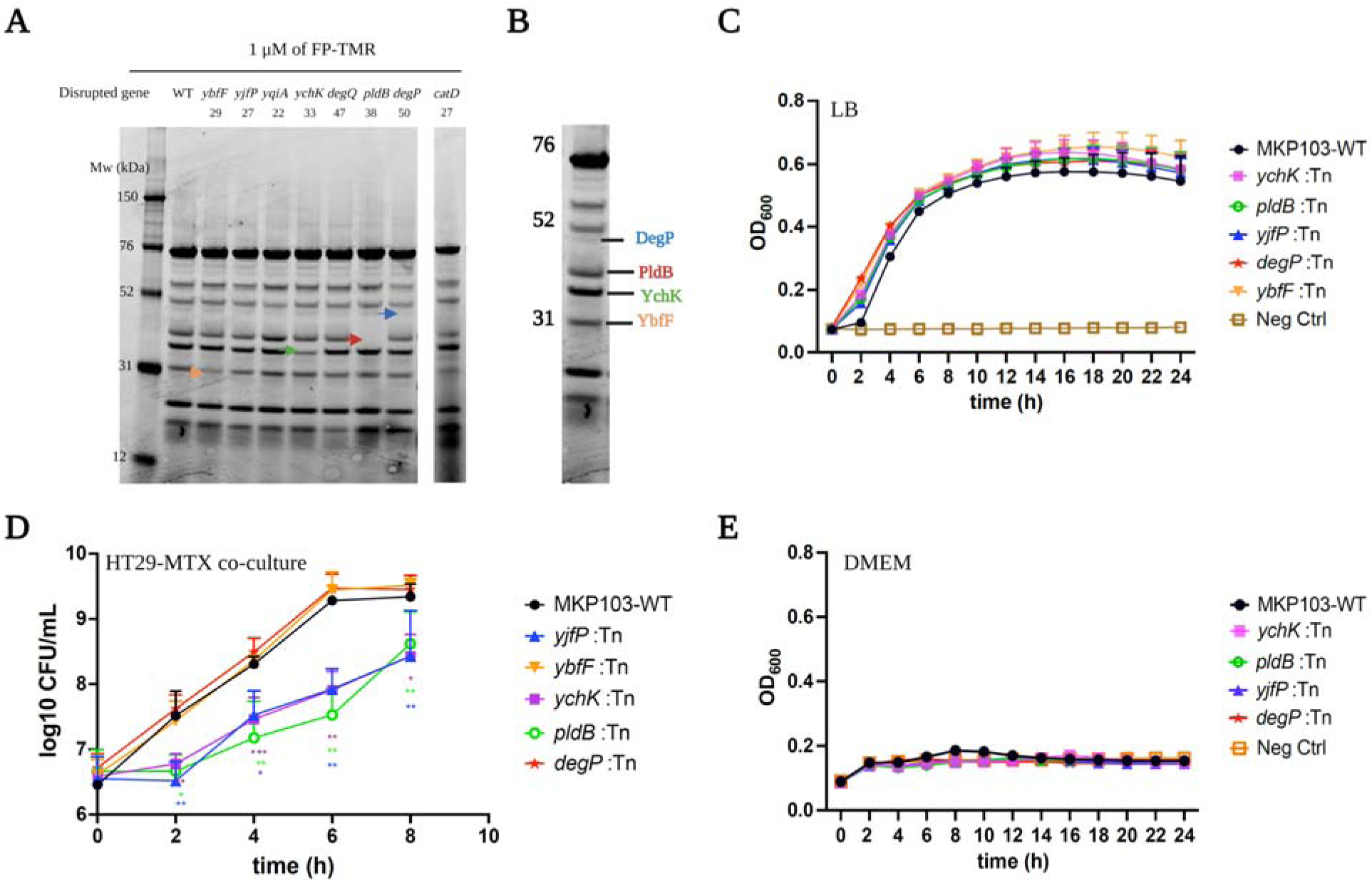
Functional validation of *K. pneumoniae* transposon mutant strains. **A**) FP-TMR labeling profiles of *K. pneumoniae* transposon mutant strains with insertions in the indicated serine hydrolase genes. Live cells were labelled with FP-TMR and lysed prior to SDS-PAGE analysis and fluorescence scanning. Arrowheads indicate labeled proteins disappearing in individual mutant strains (Red arrowhead: PldB, blue arrowhead: DegP, gree arrowhead: YchK, and orange arrowhead: YbfF). The experiment was performed three times with similar results. B) Annotated FP-TAMRA profile of *K. pneumoniae* indicating four identified probe targets. **C)** Growth curves of *K. pneumoniae* WT and SH-deficient transposon mutants in LB media, n=3 biologically independent samples. **D)** Analysis of bacterial growth in HT29-MTX co-culture model. The graph shows bacterial CFUs (log10 CFU/mL) at different time points (0, 2, 4, 6, and 8 hours) after adding *K. pneumoniae* WT and SH-deficient transposon mutants to HT29-MTX cells at an MOI of 20. Statistical significance was assessed using two-way ANOVA test with post hoc Dunnet’’s multiple comparisons tests compared with wild type (*p < 0.05, **p < 0.01, ***p < 0.001). **E**) Growth curves of *K. pneumoniae* WT and SH-deficient transposon mutants in DMEM media without FBS. Growth curves in B-E show means ± standard deviation of n=3 independent biological culture replicates.

As a first step, we investigated the growth of the selected transposon mutants in LB, a rich liquid media and found that *ych*K, *pld*B, *yjf*P, *deg*P and *ybf*F were all dispensable for growth (**Fig. 2C**). Since the primary niche for *K. pneumoniae* as a commensal is the human gut, we next evaluated growth fitness using *in vitro* models mimicking the host-interface of the gut. We adopted a co-culture model with HT29-MTX cells, a mucus-rich goblet cell line^49^. During an 8 h co-culture period with HT29-MTX cells in DMEM/10% HI-FBS, three of the tested transposon mutants (*pldB*, *ychK*, *yjfP*), showed a notable decrease in fitness compared to the WT MKP103, *deg*P:Tn and *ybf*F:Tn strains (**Fig. 2D**). This defect was evident both as a delayed onset of replication in the co-culture model and a reduced growth rate. In comparison, neither WT MKP103, nor any of the Tn-mutants was able to grow in DMEM (**Fig. 2E**), suggesting that interactions with the host cells are crucial to sustain growth in this model and may be key for understanding these phenotypes.

### PldB, YchK, and YjfP are important for the initial stages of infection in the gut

To determine whether the lysophospholipase (PldB), patatin-like lipase (YchK), and putative esterase (YjfP), play important roles in host-pathogen interactions in the gut, we used a more physiologically relevant model system that better accounts for the complexity of human colonic epithelia. This model is a 2D-co-culture model using a colonic organoid monolayer derived from human adult tissue stem cell-derived organoids using a previously published protocol ^38^ (**Fig. 3A**). Unlike 3D organoids which have enclosed apical sides, the monolayered organoids directly exposed their apical sides to the culture medium, facilitating direct interactions between the epithelial surface and the administered bacteria^50,51^. Cellular differentiation was assessed by immunofluorescence microscopy and confirmed the presence of the main cell types characteristic of mature/differentiated human colonic epithelium: mucin-1, mucin-2, and mucin-13 cells, (identified by anti-mucin antibodies, note that mucin-13 was detected with a polyclonal antiserum that might also stain other mucins) (**Fig. 3B**), enterocytes (recognized by anti-villin) and stem cells (anti-sox9 antibody). We also detected lysozyme C (by anti-lysozyme C antibody) that is produced by Paneth cells, even though this cell type is usually present in the small intestine^52^ (**Fig. 3C**).

**Figure 3.**
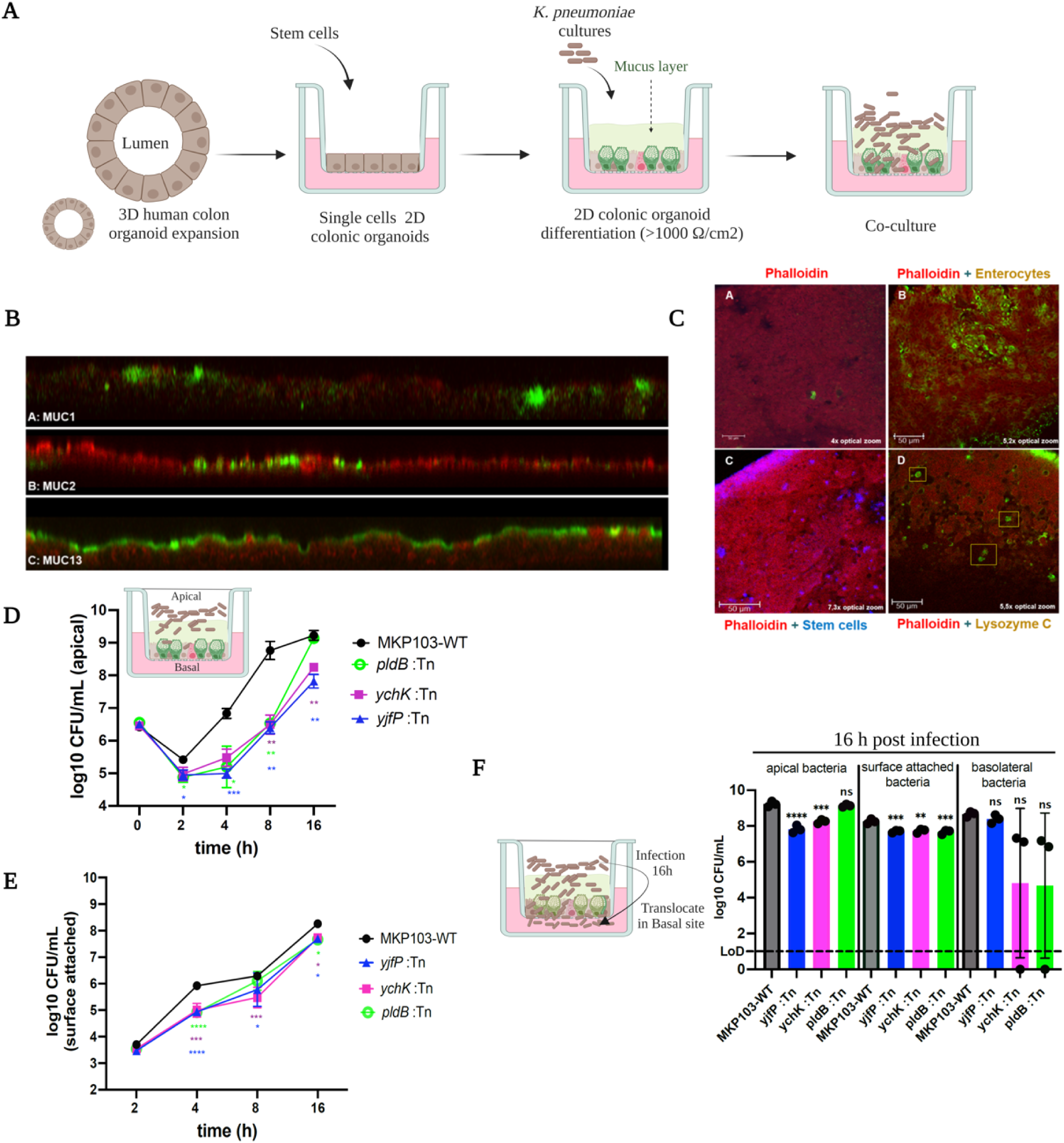
Assessment of SH-deficient *K. pneumoniae* mutants in 2D colonic organoid co-culture model. **A**) Schematic representation of the establishment and differentiation of 2D organoid cultures, including co-culturin with *K. pneumoniae* cultures. **B)** Sideview of Methacarn fixed organoid stained with Wheat Germ Agglutinin conjugated to Alexa fluorTM647 and anti-MUC antibodies combined with Alexa fluorTM488goat anti-mouse or Alexa fluorTM488 goat anti-rabbit. WGA (red) labels the cell layer as well as the mucus layer. (A) Anti-MUC1 (green) binds to the MUC1 mucin, present in the organoid samples in a relatively low abundance. The side view shows co-localization of WGA and MUC1, mainly visible at the ends of the picture. (B) Anti-MUC2 and WGA co-localize, visible in the middle part of the side view. (C) Anti-MUC13 (green) binds to the MUC13 mucin present in these organoid samples. WGA and anti-MUC13 co-localize, visible primarily on the right end of the image, where the red and green signal are at the same height. 40x magnification. **C)** Methacarn-fixed organoid cultures stained with antibodies against villin, sox9 and lysozyme C. Scale bar to represent size. (A) Negative control stained with only Phalloidin-iFluor 633 (red), Alexa fluorTM488 goat anti-mouse and Alexa fluorTM488 goat anti-rabbit (green) and Alexa fluorTM405 goat anti-rabbit (blue). 10x magnification, 4x optical zoom. (B) Organoids stained with Phalloidin-iFluor 633 to visualize the F-actin filaments of the cells (red) and anti-villin with Alexa fluorTM 488 goat anti-mouse(green) to stain villin cells. 10x magnification, 5,2x optical zoom. (C) Organoids stained with Phalloidin-iFluor 633 to visualize the F-actin filaments of the cells (red) and anti-sox9 with Alexa fluorTM 405goat anti-rabbit (blue) to stain stem cells. 10x magnification, 7,3x optical zoom. (D) Organoids stained with Phalloidin-iFluor 633 to visualize the F-actin filaments of the cells (red) and DAKOA099 with Alexa fluorTM 488 goat anti-rabbit (green) to stain lysozyme C which is secreted by Paneth cells. Yellow boxes show individual Paneth cells. 10x magnification, 5,5x optical zoom. **D)** Bacterial counts (log10 CFU/mL) in the apical part of organoid co-cultures measured at 0, 2, 4, 8, and 16 h following incubation with *K. pneumoniae* WT and SH-deficient mutants on a 2D organoid monolayer. A two-way ANOVA test with Dunnett’s multiple comparisons test was used to compare with the wild type (*p < 0.05, **p < 0.01, ***p < 0.001). **E)** Bacterial counts (log10 CFU/mL) for surface-attached bacteria to the organoid co-cultures, were measured at 0, 2, 4, 8 and 16 h following incubation with *K. pneumoniae* WT and SH-deficient mutants on a 2D organoid monolayer. A two-way ANOVA test with Dunnett’s multiple comparisons test was used, comparing mutants to the wild type, with significance levels noted as *p < 0.05, **p < 0.01, ***p < 0.001, ****p < 0.0001. **F)** *K. pneumoniae* WT and SH-deficient mutants counts (log10 CFU/mL) in the apical part, surface-attached and the basolateral compartments 16 h post infection. LoD is the lower limit of detection level. Dunnett’s one-way ANOVA was used for comparison to the WT (**p < 0.01, ***p < 0.001, ****p < 0.0001, ns indicates no significant difference).

To develop a colonization/infection model of *K. pneumoniae* in these 2D-colonic organoids, we added bacteria to the apical side of the monolayers and assessed bacterial growth by determining CFU numbers over time. We observed that the CFU counts of WT bacteria first dropped by an order of magnitude after 2 hours of co-culture compared to the inoculum. This finding could be due to the bactericidal activities of organoid-derived antimicrobial peptides. After 2 h WT bacteria started growing exponentially and approached a plateau at 8-16 hours (**Fig. 3D**). Since the bacteria were not able to grow in organoid media alone (**Supplementary figure S1**), we assume that access to host-cell derived nutrients is necessary to sustain bacterial growth in the co-culture model. After 16 h of co-culture, bacteria have translocated to the basal side of the transwell chamber suggesting damage of the epithelial barrier (**Fig. 3F**).

Upon co-culture with the organoid monolayer, the *pldB*:Tn, *ychK*:Tn, and *yjfP*:Tn mutants showed fitness defects compared to WT bacteria. The initial drop in CFU at 2 h is more pronounced in *pld*B:Tn, *ych*K:Tn and *yjf*P:Tn cells compared to WT (**Fig. 3D**) Furthermore, we observed that the outgrowth of these transposon mutant cells in the culture supernatant was delayed, but once they have resumed growth, the overall rate of growth of the transposon mutants was similar to that of WT cells. (**Fig. 3D**). From 4 h of co-culture onwards, we also observed a lower number of bacteria attached to the organoid monolayer for the three tested mutants compared to WT (**Fig. 3E**). After 16 hours of culturing, the *yjfP*:Tn mutants showed similar numbers of bacteria at the basal side as the WT despite reduced number of bacteria at the apical end. For *pldB*:Tn and *ychK*:Tn strains there was a trend towards lower bacterial numbers in the basal compartment that may be indicative of reduced or delayed translocation (**Fig. 3F**).

### Sensitivity of serine hydrolases-deficient transposon mutants to cell envelope stressors

We hypothesized that the reduced CFU levels observed in the mutants compared to WT (**Fig. 3**) could be explained by an increased sensitivity to the organoid-produced antimicrobial peptides (AMPs). To test this hypothesis, we evaluated the susceptibility of the different mutant strains to polymyxin B, a membrane-destabilizing AMP that interacts with lipopolysaccharide (LPS) ^53^. We observed an increased susceptibility to polymyxin B for both *pld*B and *yjf*P as well as for the *deg*P transposon mutant, whereas the mutant deficient in the secreted patatin-like phospholipase *ych*K had a similar susceptibility to the WT (**Fig. 4A,B**).

**Figure 4.**
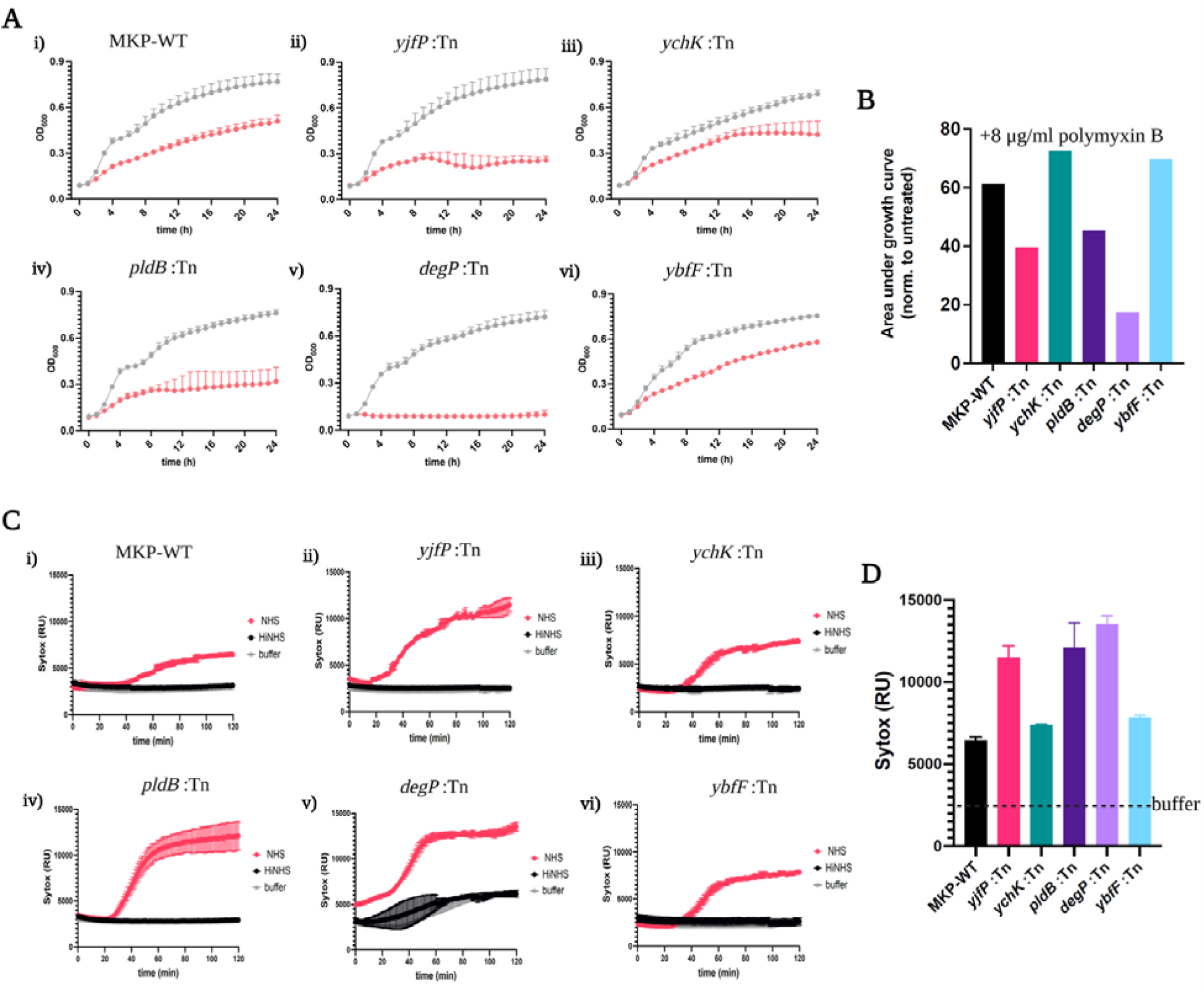
PldB, YjfP, and DegP confers resistance to envelope stressor polymyxin B and to complement killing. **A**) The growth of *K. pneumoniae* strains in absence (grey) or presence of 8 mg/ml polymyxin B (red) (mean□±□s.e.m.; n□=□5 independent replicates). B) Quantification of the area under the growth curve for treatment with polymyxin B was normalized to the growth curve of untreated cultures. **C)** Inner membrane permeabilization was evaluated in *K. pneumoniae* WT and its transposon mutants exposed to either 10% normal human serum (NHS) or 10% heat-inactivated NHS (HiNHS). The bacteria were incubated at 37°C with 1 µM SYTOX Green nucleic acid stain and SYTOX fluorescence intensity was monitored at 1 min intervals for 120 mi using a microplate reader. The presented data are the mean ± standard deviation from two independent experiments. **D)** SYTOX Green fluorescence intensity, measured after 120 min of exposure to 10% NHS as described in C. The data are presented as mean ± standard deviate on from two independent experiments. The dashed line refers t buffer permeabilization signal (RPMI + SytoxGreen).

In the Gram-negative cell envelope, LPS is not the only the target of AMPs like polymyxin, but it also plays a crucial role in resistance to complement-mediated killing^54^. A *deg*P mutant was previously reported to be more sensitive to complement-mediated killing than the parent strain^55^. To test the effects of complement mediated killing, we treated each of the mutant lines with serum and heat-inactivated serum in which components of complement are inactivated. We found that serum induced higher killing effect in the *pld*B:Tn, *yjf*P:Tn and *deg*P:Tn mutants compared to control or treatment with heat killed serum (**Fig. 4C, D)**. These results suggest a role of YjfP and PldB in shaping envelope integrity, which in turn may affect susceptibility to complement and antimicrobial peptides.

### Loss of PldB, YchK, and YjfP reduced *K. pneumoniae* virulence in *G. mellonella* larvae

The *Galleria mellonella* infection model (**Fig. 5A**) is a simple, cost-effective and easily accessible animal model that allows us to assess the effects of serine hydrolases on virulence ^56,57^. We found that upon infection with 10^5^ CFU WT *K. pneumoniae*, approximately 75% of *G. mellonella* larvae died within 2 days, whereas larvae infected with the *pld*B:Tn, *ychK*:Tn, and *yjfP*:Tn strains showed significantly higher survival rates (pL<L0.0001) (**Fig. 5B**) and approx. 65% of the larvae remained alive even after 5 days. The *ybfF*:Tn and y*qiA*:Tn mutants showed similar virulence as the WT *K.pneumoniae* MKP103 whereas, the *degP*:Tn mutant displayed reduced virulence compared to WT.

**Figure 5.**
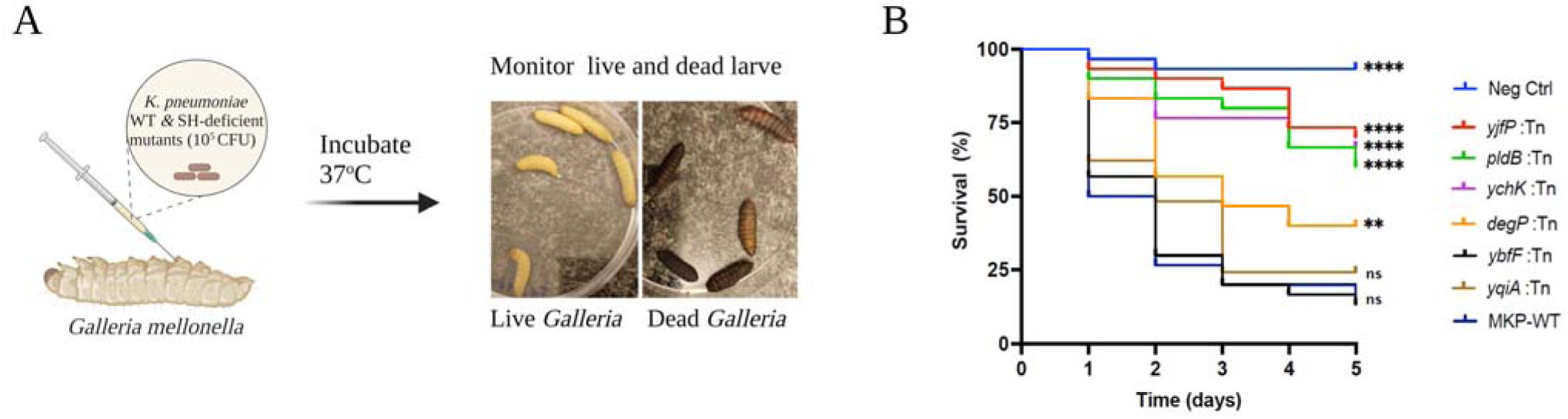
Infection of *G. mellonella* with *K. pneumoniae* strains. **A**) Schematic diagram illustrating the experimental process: Groups of *G. mellonella* larvae were injected with *K. pneumoniae* WT or SH-deficient transposon mutants and incubated at 37 °C for up to 5 days. Bacterial virulence was evaluated by monitoring of liv and dead larvae. **B)** Kaplan–Meier (KM) survival plots of *G. mellonella* larvae after inoculation of the *K. pneumoniae* WT and SH-deficient mutants. Plots show an average of 3 independent experiments with 10 larvae per group with mortality monitored daily for 5 day (N= 210). Larvae injected with PBS were used as negative control. Mutants displaying a significant difference in survival compared to the WT were determined by the log-rank (Mantel–Cox) test and are denoted (**** p < 0.0001, ** p < 0.05, ns = not significantly different).

However, phenotypic results achieved by testing of the transposon mutants might be confounded by secondary mutations or polar effects. To rule out secondary mutations that may account for these phenotypes, we retrieved additional mutants from the Manoil *K. pneumoniae* 3-allele transposon mutant library^45^ representing individual clones where the transposons were localized within the same gene, but at a different location. Assessment of the virulence of different transposon mutants with insertions in *pld*B (n=3 available strains), *yjf*P (n=3) and *ych*K (n=2) (**Supplementary Figure S2**) all showed similar phenotypes, suggesting that transposon insertion in the respective gene accounts for the observed phenotype rather than secondary mutations. In conclusion, assessment of virulence of the *ych*K, *pld*B, and *yjf*P mutants supports a role of these enzymes in virulence of *K. pneumoniae*.

### Biochemical and structure characteristics of YjfP and other serine hydrolases

Of the seven uncharacterized serine hydrolases that we identified in our ABPP proteomics studies, we were able to purify YjfP, YqiA, and PldB (**Supplementary figure S3**). To assess the substrate preference of recombinantly expressed and purified YjfP, YqiA, and PldB, we examined their ability to cleave various commercially available fluorogenic substrates (**Fig. 6A, B, C**). Our results indicated that YjfP effectively hydrolyzes saturated lipid esters with chain lengths from C2 to C4, showing a strong preference for the short-chain C2 acetate substrate (**Fig. 6A**). YqiA cleaved a variety of artificial substrates ranging from C2 to C10, with the highest activity observed for the C7 heptanoate substrate (**Fig. 6B**). PldB hydrolyzed saturated lipid esters ranging in chain length from C2 to C8, showing a particularly strong preference for the C2 and C4 chain lengths (**Fig. 6B**).

**Figure 6.**
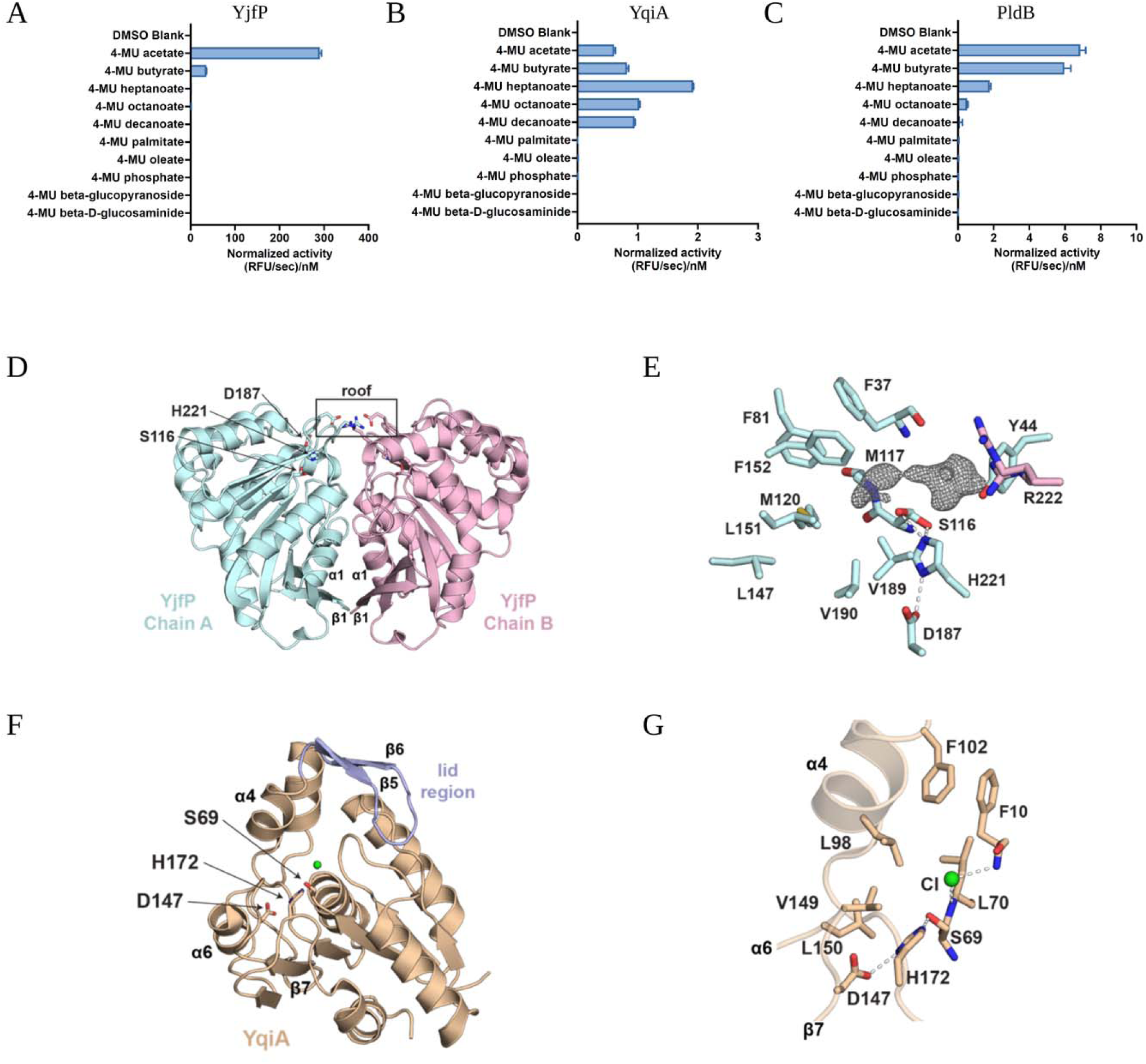
Biochemical characterization of YjfP, YqiA, and PldB and structure of hydrolases YjfP and YqiA. Normalized cleavage rate (, in relative fluorescent unit (RFU)/s per nM of protein) of 4-methylumbelliferone (4-MU) substrates by YjfP (**A**), by YqiA (**B**), and by PldB. (**C**). The presented bars represent the mean ± standar deviation (n = 3 technical replicates). **D**) YjfP crystallised as a dimer (cyan and pink), where the dimer interface is mediated by connections across β strands 1 and α helices 1 from each chain. Two salt bridges are formed at the to of the dimer, creating a roof above a putative substrate binding cavity. This cavity contains the catalytic triad; Ser116, Asp187, and His221. **E**) Interactions are shown between side chains of the catalytic triad (modelled in two conformations). Displayed around Ser116 is the mFo-DFc map contoured at 3.5 for a putative ligand. This putativ ligand likely interacts with a hydrophilic portion of the active site (Tyr44, Ser116, and Arg222 modelled in tw conformations). Part of the density is in the oxyanion hole, coordinated by backbone amides from Met117 an Phe37. Beyond the oxyanion hole, several hydrophobic side chains are shown. **F**) YqiA (beige) crystallised as a monomer, where β strands 4 and 5 form a lid region. The catalytic triad (Ser69, Asp147, His172) is more expose than in the YjfP structure. **G**) Interactions between side chains of the catalytic triad are shown. A chlorine atom (green) is modelled in the oxyanion hole coordinated by backbone amides of Leu70 and Phe10. Several hydrophobic side residues on α helix 4 and a loop between β strand 7 and α helix 6 are likely important for substrate recognition.

We also determined the crystal structures of YjfP and YqiA to 1.3 Å and 1.5 Å, respectively (**Fig. 6D, E, F, G, Suppl. Tables S4 and S5, Suppl. Figures S5, S6**), but we were unable to crystalize purified PldB protein. YjfP has the canonical α/β-hydrolase fold which is typical of serine hydrolases^58^, with a catalytic triad conserved at residues Ser116, Asp187, and His221 (**Fig. 6D).** An 8-stranded β-sheet is central to the α/β-hydrolase fold, sandwiched between α-helices 1 and 8, and α-helices 2-7 **(Supplementary Figure S5).** Serine hydrolases often contain a flexible lid region between α-helices 4 and 5 that mediates substrate specificity^59^. However, this is absent in YjfP, which instead has a short loop connecting α-helices 4 and 5. Consistent with our gel filtration results (**Supplementary Figure S4**), the asymmetric unit of the YjfP crystal contains a homodimer that creates a closed cavity around the catalytic triad. Submission of the YjfP structure to PISA, an online tool to analyse protein interfaces, predicts the dimer interface to be biologically relevant, scoring a maximum CSS of 1.0^60^. The dimer interface covers 11 % of each protein surface area and is mediated mostly through β-strand 1 and α-helix 1. At the top of this interface, Asp188 and Arg220 from one chain form salt bridges with Arg220 and Asp188 from the second chain. These residues are on the same loops as Asp187 and His221 from the catalytic triad, forming a roof above the putative substrate binding site. Inside the substrate binding cavity there is discontinuous density, potentially corresponding to a cleaved substrate that may have co-purified (**Fig. 6E**). This density approaches a hydrophobic face within the cavity that would not accommodate longer carbon chains than butyrate, explaining the sharp drop-off in activity for C7 and longer chain substrates (**Fig. 6A**).

Similar to YjfP, YqiA has a canonical α/β hydrolase fold^58^, with a conserved catalytic triad of residues Ser69, Asp147, and His172 (**Fig. 6F**). However, there are several features in YqiA that differ from YjfP. Firstly, the asymmetric unit contains a monomer, consistent with gel filtration results (**Supplementary Figure S4**), and PISA predicts there are no crystal contacts that are likely to create a dimer^60^. Unlike YjfP, the YqiA active site is much more open despite the presence of a small lid region. Within the oxyanion hole there is a chlorine atom and a hydrophobic face is formed around the active site (**Fig. 6G**). This hydrophobic, open face likely directs substrate specificity towards hydrophobic carbon chains. Since the lid region is small and removed from the active site, it might not accommodate the 4MU leaving group of the fluorogenic substrates well, which might explain the much lower activity rates of YqiA compared to YjfP.

Together, the substrate profiling and structural characterization of YjfP and YqiA suggest they are short-chain esterases. the YjfP structure showed that an acetate group would fit well into the small hydrophobic pocket near the active site Ser116, whereas chains longer than butyrate would not, thus giving an explanation for the observed substrate selectivity (**Fig. 6A)**. Our data suggest that YjfP may function as a deacetylase.

## Discussion

Serine hydrolases play important roles in various biological processes of both bacteria and host cells yet our understanding of these enzymes in *K. pneumoniae* remains limited. Using a chemical proteomic approach, we have identified 10 previously uncharacterized serine hydrolases. Using transposon mutants of the hydrolases PldB, YchK, and YjfP we observed growth defects in co-culture models with HT29-MTX cells, human colonic organoids and in a *Galleria mellonella* infection model, suggesting putative roles in both gut colonization and infection. In the *G. mellonella* infection model, alternative clones with transposon insertion in the same gene displayed the same phenotype suggesting its specificity.

Based on previous annotation and bioinformatic analysis, the previously uncharacterized PldB, is a lysophospholipase L2 located at the cytoplasmic membrane. YjfP is annotated as a putatively cytosolic esterase. YchK which possesses a patatin-like phospholipase domain is predicted to be secreted. Of note, despite similarities of the phenotypes in the complex organoid and *G. mellonella* infection models, we observed some important differences that might give clues about the underlying molecular functions of these enzymes. Both the *pld*B and *yjf*P-deficient strains showed an increased susceptibility to AMPs, whereas the *ych*K mutant did not.

AMPs are short amphiphilic peptides that, in general, act by disrupting bacterial membranes. Polymyxin B was shown to exert its membrane-destabilizing effect by binding to the lipid A anchor of LPS in the outer membrane of Gram-negative bacteria^61^. AMPs produced by human cells, e.g., the defensins, are considered to act by similar mechanisms. In the human gut, the specialized secretory Paneth cells in the small intestine produce alpha-defensins (as reviewed by Bevins and Salzman^62^), whereas epithelial cells contribute through production of beta-defensins (see review by Gallo and Hooper^63^). In our colonic stem-cell derived organoid model, the presence of Paneth cells was indirectly detected, and we therefore assume that bacteria in the organoid-model are exposed to both alpha– and beta-defensins. Human beta-defensin 1 is expressed by HT29 cells^64^ and thus may affect bacterial growth in the HT29-MTX co-culture model. Of note, AMPs are also important effectors of insect innate immunity (as reviewed by Stacek^65^), and *G. mellonella* has been shown to mount pathogen-specific AMP responses upon infection with diverse bacteria and fungi^66^. Thus, an increased susceptibility to AMPs could be detected in both organoid and HT29-MTX co-culture models as well as in the *G. mellonella* infection model.

The most pronounced increase in polymyxin B-sensitivity was observed for the *deg*P-mutant, which is consistent with a previous study^27^. This mutant, however, did not show any fitness defect in the HT29-MTX model (and was hence not tested in organoids) and the reduction in virulence in the *G. mellonella* model was less pronounced (and less stable across different clones) compared to the *pld*B and *yjf*P mutants. This discrepancy could be explained by differences in the susceptibility to polymyxin B compared to the AMPs that are endogenously produced in the co-culture and infection models. However, this outcome might also indicate that a reduced susceptibility to AMPs might contribute to the phenotypes in the co-culture and infection models, but on its own is not sufficient to explain them. It is possible that the putative deficiency in cell envelope integrity in the *pld*B and *yjf*P mutants leads to multiple downstream effects that collectively contribute to reduced virulence and fitness.

We can only speculate about by which mechanism PldB and YjfP may affect cell envelope integrity, but roles in membrane remodeling, tailoring modifications of LPS cell wall components, or posttranslational modifications of proteins that are involved in cell wall and LPS synthesis could be plausible. Of note, secondary acylation of LPS in *K. pneumoniae* affects susceptibility to polymyxin B and several other AMPs^67^, suggesting that remodeling of the lipid A anchor through previously unidentified hydrolases or transferases could affect cell membrane integrity and AMP susceptibility. However, cleavage of lipid substrates with longer chain fatty acid esters does not appear a likely physiological function of YjfP. The narrow substrate profile and structural characterization of this enzyme suggest that is a deacetylase, that may act on a small molecule, peptide or protein. PldB, in contrast accepts a broader range of substrates cleaving synthetic fluorogenic substrates up to C8 chain length. The interesting phenotypes associated with these SH mutants and the architecture of the YjfP (and YqiA) active sites, warrant further investigation into the native substrates of each enzyme.

The *pld*B and *yjf*P mutant also showed increased sensitivity to complement killing. Since complement activation involves pore formation in the outer membrane and complement resistance is mediated by the protective effects of LPS or capsule polysaccharide ^68^, these results support the notion that PldB and YjfP might also be partially responsible for the modulation of LPS production and/or constructing the outer membrane. Recently, another chemoproteomic study identified BT4193, a homolog of human dipeptidyl peptidase 4, in the Gram-negative gut commensal *Bacteroides thetaiotaomicron* ^23^. Due to its periplasmic location, the authors suspected a role in cell envelope integrity and observed that a BT4193 mutant displayed increased sensitivity for killing by both vancomycin and polymyxin B. However, treatment of WT cells with a BT4193 inhibitor did not phenocopy these deficiencies, so the authors concluded that the phenotype of the ΔBT4193 mutant might be caused by a structural role of this protein rather than by its enzymatic function^23^. Whether the putative role of PldB and YjfP on AMP and complement susceptibility will be due to its enzymatic functions as a lipases or esterase or due to structural roles, remains to be determined.

In contrast to the cell-membrane associated putative lysophospholipase PldB and putatively cytosolic deacetylase YjfP, the putative phospholipase YchK is predicted to be secreted. As may be expected for a secreted phospholipase, the YchK-mutant did not show any phenotype related to susceptibility to AMP and envelope integrity. Secreted lipases have been described as virulence factors for a number of bacterial pathogens and their hydrolytic function mostly associated with the destruction of defense barrier and toxicity ^69–71^. One interesting observation that we made is that *K. pneumoniae* is unable to grow in organoid media alone or in the DMEM medium alone used in the HT29-MTX culture. This means that to sustain growth in the organoid co-culture model, bacteria need to access nutrients from the host cell layers. Both the colon organoid and HT29-MTX monolayer produce a pronounced mucus layer. In the gut, the mucus layer, which is predominantly composed of glycoproteins, is an essential part of human colonic epithelia and acts as the primary defense against bacteria residing in the colon^72–74^. In the human gut, the mucus layer of colonic epithelia has a surface layer of phosphatidylcholine (PC) which contributes to barrier function and immune regulation ^75^. Bacterial phospholipases in this layer can convert PC into lyso-PC, thereby disturbing the integrity and protective function of the mucus structure and enabling damage to epithelial cells ^75–77^. For the gastric pathogen *H. pylori* secreted lipases are responsible for breaching a mucus-associated defensive phospholipid layer, allowing the pathogen to partially invade the gastric mucus ^78,79^. A similar role is plausible for the putatively secreted patatin-like phospholipase YchK of *K. pneumoniae*. Its hydrolytic activity could enable either direct access to phospholipid-derived nutrients or contributing to the break-down of defense barriers that might facilitate cell invasion and extraction of nutrients. However, the origin of the surface layer of PC in the human colon is unclear (Review by ^75^) and it might be derived from the ileum or jejunum. It thus remains to be determined if a similar PC layer is in fact found in our organoid model. An alternative role for YchK at the host-pathogen interface in the organoid model as well as in *Galleria infection* model could be that it directly acts on host cell membranes.

Our results indicate that PldB, YjfP and YchK may be promising targets for further evaluation as anti-virulence targets. One of the last-resort antibiotics for treatment of infections with carbapenemase producing *K. pneumoniae* is the AMP colistin^80^. The emergence of colistin-resistance in the clinic is therefore of great concern^81^ and strategies to reverse colistin resistance are being explored^82^. The sensitization to AMPs observed for *pld*B, *yjf*P and *deg*P mutants is a significant finding and we suggest that inhibitors of either of these enzymes should be tested for synergy with colistin and other AMP antibiotics.

Another potential drug target that was identified in our dataset but was not validated further, is the carboxylesterase *Bio*H. This enzyme has a putative role in the biosynthesis of biotin^48^ and the absence of viable mutants within the Manoil Lab transposon mutant library^45^ suggests that it might be essential. Of note, in *Mycobacterium tuberculosis,* biotin metabolism is already considered a viable antibacterial drug target pathway^83,84^, and efforts to target its three bioH isoenzymes are underway^85^.

For ABPP, enzyme targets are identified through covalent interaction with a small molecule probe, thus focusing the set of targets that are often likely to be druggable and provides direct assays for development of target-specific inhibitors. There is an increasing pool of serine-hydrolase-reactive chemotypes that continue to produce target-specific inhibitors that can be used as tools to further investigate biological function while also sometime serving as antimicrobial drug candidates. The structural information of YjfP provided in this work may enable structure-based design of specific YjfP inhibitors. Alternatively, inhibitors may be identified through traditional target-based screening approaches^44^, or through competitive ABPP which simultaneously assesses the selectivity profile of candidate inhibitors against all of the active serine hydrolase targets in a sample^17,20,86–88^. Importantly, if performed on live cells, as in the current study, cell-based competitive ABPP, can also confirm the accessibility of the targets to small molecule inhibitors^17,86^.

We believe that further functional characterizations of the uncharacterized serine hydrolases identified will significantly enhance our understanding of *K. pneumoniae* pathogenesis and colonization and provide the basis for their validation as anti-virulence and antibacterial drug candidates that are urgently needed in times of emerging multidrug resistance of this critical priority pathogen.

## Acknowledgements

This work was funded by a Centre for New Antibacterial Strategies (CANS) starting-grant through the Trond-Mohn Foundation to C.S.L. We are grateful for additional financial support through the Aurora Outstanding Career Development program at UiT through C.S.L. The research stay of M. J. U. at UMC Utrecht was generously supported by The National Graduate School in Infection Biology and Antimicrobials (IBA). LC-MS/MS analyses were carried out at the UiT Proteomics and Metabolomics Core Facility (PRiME). The PRiME is part of the National Network of Advanced Proteomics Infrastructure (NAPI), funded by the Research Council of Norway INFRASTRUKTUR-program (project number: 295910). We also thank Fernanda Paganelli for her support during the initial establishment of the organoid model, Janetta Top, Rob Willems and Bart W. Bardoel for helpful discussions, Coco Duizer for discussions in the lab, and Karin Strijbis for generously providing the Anti-MUC13 antibody, all from UMC Utrecht. Some of this work was supported by the New Zealand China – Maurice Wilkins Centre Collaborative Research Programme (to M.F.) and was conducted during tenure of The Sir Charles Hercus Health Research Fellowship of the Health Research Council of New Zealand (to M.F.). This research was undertaken in part using the MX2 beamline at the Australian Synchrotron, part of ANSTO, and made use of the ACRF detector. F.M. was supported by the European Union’s Horizon 2020 research program 787 H2020-EU-ITN-EJD (CORVOS #860044)

## Ethics declaration

The organoids used were obtained from the Foundation Hubrecht Organoid Biobank (Utrecht, The Netherlands) under TC-Bio protocol number 14-008 and used according to informed consent.

Human blood was isolated after informed consent was obtained from all subjects in accordance with the Declaration of Helsinki. Approval was obtained from the medical ethics committee of the UMC Utrecht, The Netherlands

## Author contributions

Conceptualization: C.S.L., M.J.U., Investigation: M.J.U., G.R., J.Z., T.U., L.v.E, P.H., F.M.; M.C.V., Data analysis: M.J.U., G.R., J.Z., T.U., L.v.E, P.H., M.C.V., M.B., M.F., M.R.d.Z., M.J., C.S.L., Supervision: C.S.L., M.J., M.R.d. Z., M.F., M. B., Funding acquisition: C.S.L., Writing – Original manuscript draft: M.J.U. and C.S.L., Writing – Editing: All authors.

