## Supplementary Information for "Activity-Based Protein Profiling Identifies *Klebsiella pneumoniae* Serine Hydrolases with Potential Roles in Host-Pathogen Interactions"

#### Table of Contents

Table S1: List of bacterial strains used in this study…………………………………………….. page 2
Table S2: List of serine hydrolases identified in *K. pneumoniae* by MS-ABPP ………………… page 3
Table S3: The homology percentage data used for Figure 1E
Table S4: Collection statistics of x ray crystallography datasets used ……………………….…….… page 3
to determine the structures of YjfP and YqiA………………………………………….... page 4
Table S5: Refinement and validation statistics of YjfP and YqiA crystal structures………………page 4

Figures S1: Growth curves of *K. pneumoniae* MKP103 and isogenic SH-deficient
transposon mutant on organoids secreted substances…………………………………..…..page 5
Figure S2. Infection of *G. mellonella* with *K. pneumoniae*………………...………………………page 6
Figure S3. Coomassie-stained images of purified YjfP, YqiA, and PldB……………………… page 7
Figure S4. Gel-filtration chromatogram of YjfP, YqiA. ……………………………….……… page 7
Figure S5. Secondary structures of YjfP……………………………………………………………page 8
Figuer S6. Secondary structures of YqiA…………………………………………...…….…page 8

Supplementary References ……………..…………………………………………..….…page 9

### Supplementary tables

**Table S1**. List of bacterial strains used in this study.

| Strain | Description | Reference/Source |
|---|---|---|
| *K. pneumoniae* MP103 | KPNIH1 derivative with the KPC-3 carbapenemase-encoding gene deleted and Parent strain of Moinul Transposon Mutant Library | [1] |
| *degP*:Tn | Transposon insertion mutant in MKP103 KPNIH1_04540 | |
| *ychK*:Tn | Transposon insertion mutant in MKP103 KPNIH1_15760 | |
| *ybfF*:Tn | Transposon insertion mutant in MKP103 KPNIH1_07725 | |
| *catD*:Tn | Transposon insertion mutant in MKP103 KPNIH1_12245 | |
| *pldB*:Tn | Transposon insertion mutant in MKP103 KPNIH1_00895 | |
| *degQ*:Tn | Transposon insertion mutant in MKP103 KPNIH1_23755 | |
| *yjfP*:Tn | Transposon insertion mutant in MKP103 KPNIH1_02230 | |

**Table S2.** List of serine hydrolases identified in *K. pneumoniae* by MS-ABPP

| **Gene** | **Previous annotation** | **MW [kDa]** | **Strain KPNIH1 locus tag** |
|---|---|---|---|
| *degP* | Serine endoprotease | 49,5 | KPNIH1_04540 |
| *ychK* | Patatin-like phospholipase | 33,3 | KPNIH1_15760 |
| *ybfF* | Acyl-CoA esterase | 28,5 | KPNIH1_07725 |
| *catD* | 3-oxoadipate enol-lactonase | 27,3 | KPNIH1_12245 |
| *pldB* | Lysophospholipase L2 | 38,2 | KPNIH1_00895 |
| *degQ* | Serine endoprotease | 47,2 | KPNIH1_23755 |
| *YqiA* | Esterase | 21,5 | KPNIH1_22780 |
| *YcfP* | Esterase | 21,1 | KPNIH1_09915 |
| *bioH* | Pimeloyl-ACP methyl ester esterase | 28,3 | KPNIH1_24570 |
| *yjfP* | Esterase | 26,5 | KPNIH1_02230 |

**Table S3**. The homology percentage data used for Figure 1E.

*All BLASTp results for a homolog of the K. pneumoniae SHs are provided in* ***Extended Dataset 2***

| | DegP | YchK | YbfF | CatD | PldB | DegQ | YqiA | YcfP | bioH | YjfP |
|---|---|---|---|---|---|---|---|---|---|---|
| *Bifidobacterium adolescentis ATCC 15703* | 0 | 0 | 0 | 0 | 0 | 0 | 0 | 0 | 0 | 28.4 |
| *Bifidobacterium longum NCC2705* | 0 | 0 | 26.6 | 24.3 | 0 | 0 | 0 | 0 | 0 | 27.4 |
| *Collinsella aerofaciens ATCC 25986* | 0 | 0 | 0 | 0 | 0 | 0 | 0 | 0 | 0 | 24.8 |
| *Bacteroides caccae ATCC 43185* | 0 | 0 | 0 | 0 | 0 | 0 | 0 | 0 | 0 | |
| *Bacteroides fragilis ATCC 25285* | 0 | 0 | 0 | 0 | 0 | 0 | 0 | 0 | 0 | 25.7 |
| *Bacteroides ovatus ATCC 8483* | 0 | 0 | 0 | 0 | 0 | 0 | 0 | 0 | 20.0 | 24.4 |
| *Parabacteroides distasonis ATCC 8503* | 0 | 0 | 0 | 0 | 0 | 0 | 0 | 0 | 0 | |
| *Prevotella copri DSM 18205* | 0 | 0 | 0 | 0 | 0 | 0 | 0 | 0 | 0 | 41.3 |
| *Clostridium sporogenes ATCC 15579* | 0 | 0 | 0 | 0 | 0 | 0 | 0 | 0 | 0 | 0 |
| *Enterococcus faecalis V583* | 0 | 0 | 26.6 | 0 | 0 | 0 | 0 | 0 | 23.6 | 0 |
| *Enterococcus faecium ATCC BAA-472* | 0 | 0 | 0 | 0 | 0 | 0 | 0 | 0 | 29.3 | 0 |
| *Lactobacillus ruminis ATCC 25644* | 0 | 0 | 0 | 0 | 0 | 0 | 0 | 0 | 0 | 0 |
| *Ruminococcus gnavus ATCC 29149* | 0 | 0 | 0 | 0 | 0 | 0 | 0 | 0 | 0 | 0 |
| *Fusobacterium nucleatum subsp. nucleatum ATCC 23726* | 0 | 0 | 0 | 0 | 0 | 0 | 0 | 0 | 0 | 0 |
| *Edwardsiella tarda ATCC 23685* | 77.9 | 0 | 0 | 0 | 0 | 62.0 | 0 | | 0 | 0 |
| *Enterobacter cancerogenus ATCC 35316* | 91.9 | 0 | 0 | 0 | 84.9 | 82.0 | 0 | 90.0 | 0 | 0 |
| *Escherichia coli K12 MG1655* | 90.2 | 81.1 | 0 | 24.7 | 79.7 | 0 | 0 | 0 | 0 | 0 |
| *Klebsiella pneumoniae subsp. pneumoniae KPNIH1* | 100.0 | 100.0 | 100.0 | 100.0 | 100.0 | 100.0 | 100.0 | 100.0 | 100.0 | 100.0 |
| *Akkermansia muciniphila ATCC BAA-835* | 0 | 0 | 0 | 0 | 0 | 0 | 0 | 0 | 0 | 0 |
| *Homo sapiens* | 0 | 0 | 0 | 0 | 25.2 | | | | 30.2 | |

**Table S4**. Collection statistics of x ray crystallography datasets used to determine the structures of YjfP and YqiA. Values in parentheses refer to the high-resolution shell.

| | **YjfP** | **YqiA** |
|---|---|---|
| Beamline | Australian Synchrotron MX2 | Australian Synchrotron MX2 |
| Wavelength (Å / keV) | 0.954 / 13.00 | 0.954 / 13.00 |
| Detector | DECTRIS EIGER X 16M | DECTRIS EIGER X 16M |
| Space group | *C* 2 | $P\ 3_2\ 2$ |
| a, b, c (Å) | 102.71 68.20 65.36 | 62.46 62.46 80.36 |
| α, β, γ (°) | 90.00 77.16 90.00 | 90.00 90.00 120 |
| Rotation range (°) | 250 | 310 |
| Resolution (Å) | 45.11-1.30 (1.32-1.30) | 44.87-1.50 (1.53-1.50) |
| Total Reflections | 513,218 (23,779) | 507,705 (23,786) |
| Unique Reflections | 106,974 (5,160) | 29,609 (1,434) |
| Multiplicity | 4.8 (4.6) | 17.1 (16.60) |
| Completeness (%) | 99.2 (97.4) | 100.00 (100.00) |
| I/σ(I) | 9.6 (1.7) | 13.8 (1.7) |
| $CC_{1/2}$ | 0.998 (0.369) | 0.999 (0.568) |
| $R_{merge}$ | 0.115 (2.702) | 0.121 (5.951) |
| $R_{pim}$ | 0.088 (2.083) | 0.043 (2.148) |

**Table S5**. Refinement and validation statistics of YjfP and YqiA crystal structures.

| | **YjfP** | **YqiA** |
|---|---|---|
| PDB ID | 9BD4 | 9BI7 |
| Resolution Range (Å) | 45.11-1.30 | 44.87-1.50 |
| Reflections, working | 101,407 | 28,024 |
| Reflections, free | 5,336 | 1,528 |
| $R_{work}$ (%) | 16.13 | 17.46 |
| $R_{free}$ (%) | 17.48 | 21.31 |
| Number of residues | 479 | 192 |
| Number of waters | 305 | 94 |
| Ligand | | 2 Ca, 1 Cl |
| Average *B* factors ($Å^2$) | 16.22 | 30.84 |
| Ligands | | 23.88 |
| Waters | 25.01 | 33.35 |
| RMSD | | |
| Bonds (Å) | 0.006 | 0.009 |
| Angles (°) | 0.89 | 1.00 |
| Ramachandran | | |
| Favoured (%) | 97.89 | 98.42 |
| Outliers (%) | 0 | 0 |
| Rotamer outliers (%) | 0.25 | 0 |
| Molprobity score | 0.88 | 1.08 |

### Supplementary figures

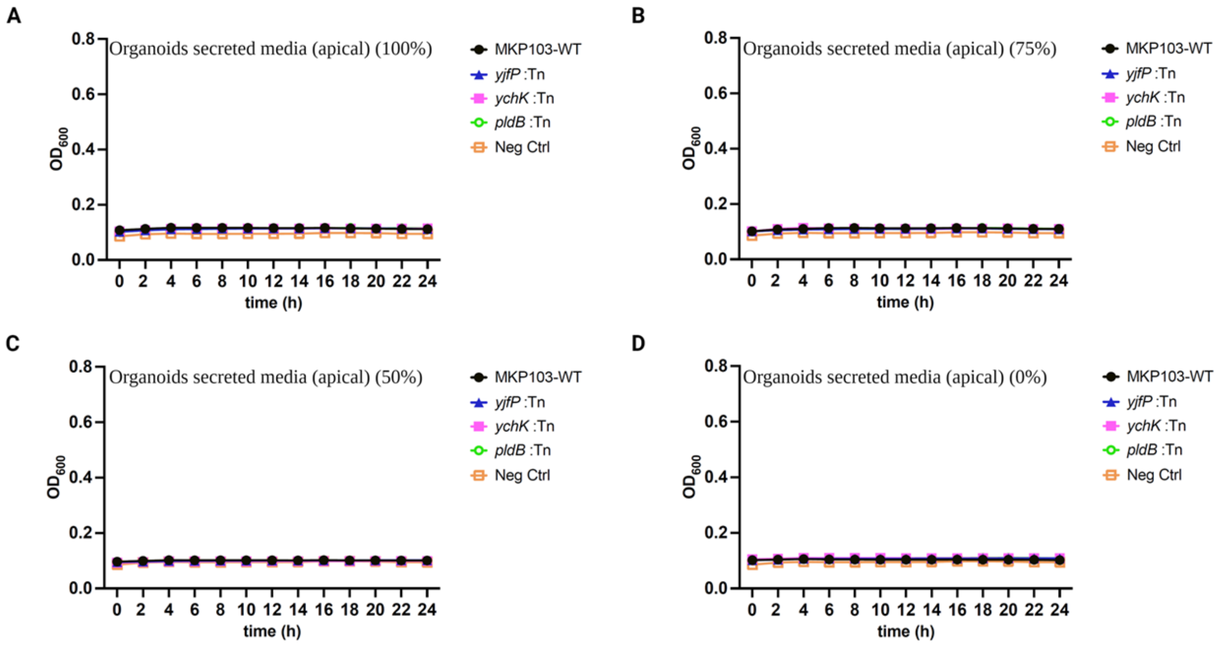


**Figure S1**. **Growth curves of *K. pneumoniae* MKP103 and isogenic SH-deficient transposon mutant on organoids secreted substances**. The growth curves of *K. pneumoniae* and SH-deficient mutants were measured over 24 hours in: **A**) media containing 100% secretions from organoid monolayers (collected from the apical side), **B**) media with 75% organoid secretions, **C**) media with 50% organoid secretions, and **D**) media without organoid secretions (consisting only of differentiated organoid media). Growth curves in A-D show means standard deviation of n=3 independent biological culture replicates

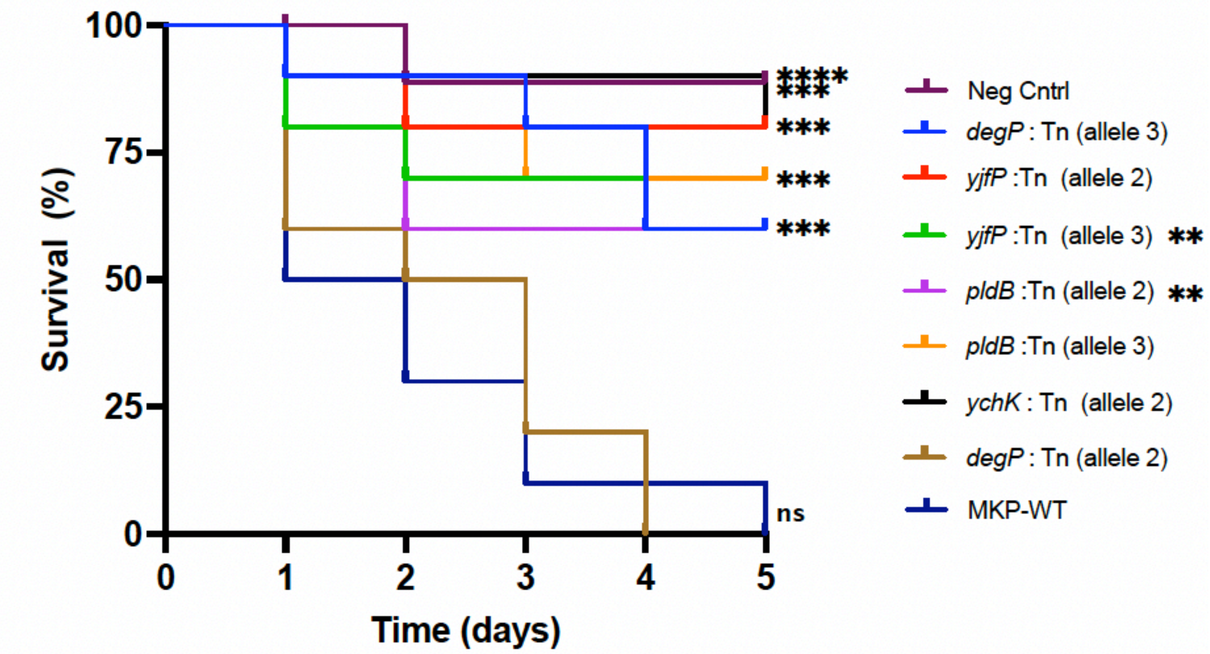


**Figure S2**. **Infection of *G. mellonella* with *K. pneumoniae*.** Kaplan–Meier (KM) survival plots of *G. mellonella* larvae after inoculation of the *K. pneumoniae* WT and SH-deficient mutants. Different transposon mutants allele displaying a significant difference in survival compared to the WT were determined by the to log-rank (Mantel–Cox) test and are denoted (**** $p < 0.0001$, **** $p < 0.001$, ** $p < 0.05$, ns = not significantly different).

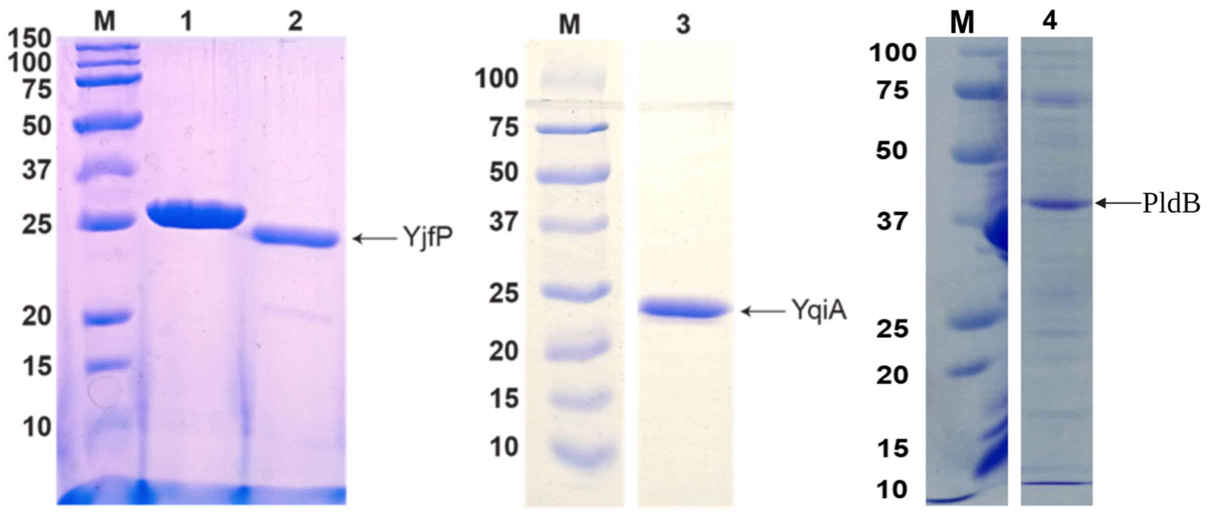


**Figure S3**. SDS PAGE of purified YjfP, YqiA, and His6-PldB. YjfP, YqiA, and PldB purity were assessed by loading onto SDS PAGE gels of 12 % polyacrylamide. Molecular weight markers are in lanes labelled M and labelled by their size in kDa. His6 YjfP was loaded into lane 1, YjfP after 3C cleavage loaded into lane 2, YqiA loaded into lane 3, and His6-PldB loaded into lane 4. The predicted masses after cleavage of YjfP, YqiA, and PldB are 27 kDa, 22 kDa, and 38 kDa, respectively.

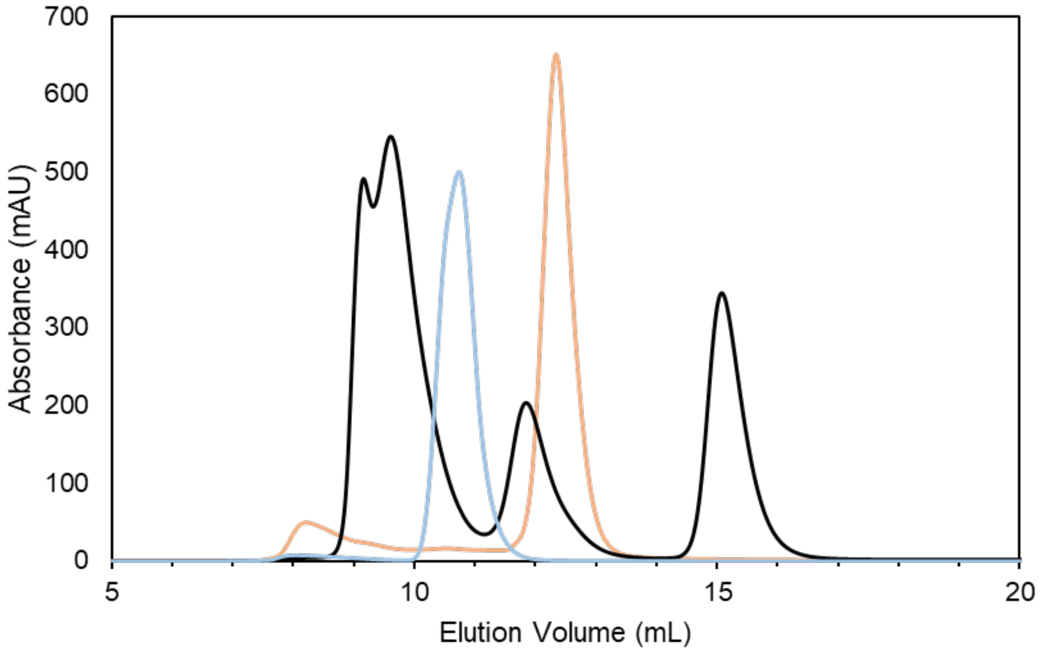


**Figure S4**. **Gel-filtration chromatogram of YjfP and YqiA**. Gel filtration of YjfP (cyan), YqiA (orange), and protein standards (black). The protein standards run include thyroglobulin (670 kDa), bovine gamma globulin (158 kDa), chicken ovalbumin (44 kDa), and equine myoglobin (17 kDa). YjfP elutes after bovine gamma globulin and before chicken ovalbumin, consistent with a 54 kDa homodimer. YqiA elutes after chicken ovalbumin consistent with a 22 kDa monomer.

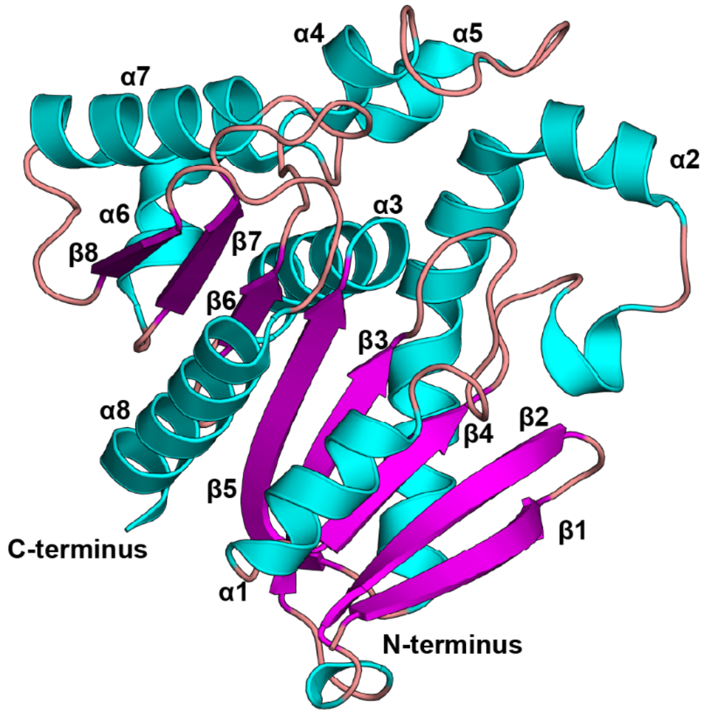


**Figure S5**. **Secondary structures of YjfP**. Secondary structure elements are displayed as follows: α helices (cyan), β sheets (magenta), and loops (beige).

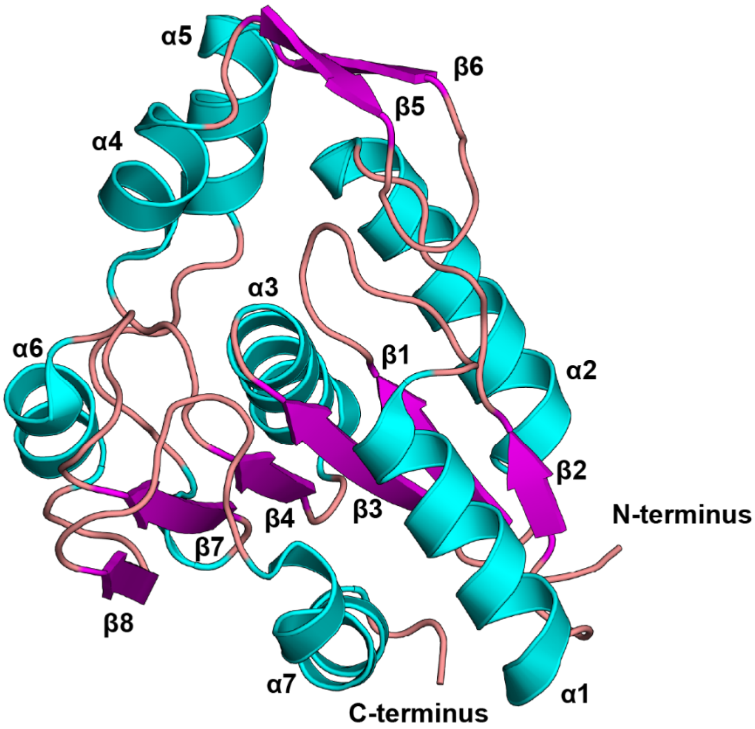


**Figure S6**. **Secondary structures of YqiA.** Secondary structure elements are displayed as follows: α helices (cyan), β sheets (magenta), and loops (beige).
